# A comparative analysis of promoter-proximal pausing reveals kinetic and distributional dimensions of variation

**DOI:** 10.64898/2026.06.01.729264

**Authors:** Xin Zeng, Gilad Barshad, Rebecca Hassett, Edward J Rice, Charles G Danko, Adam Siepel, Yixin Zhao

## Abstract

Promoter-proximal pausing of RNA polymerase II is a key regulatory checkpoint in metazoan transcription. Despite extensive study of this process, quantitative methods for comparing pausing dynamics across biological contexts have been lacking. Here we introduce a model-based framework for rigorous comparative analysis of both pause-escape kinetics and pause-site distributions. We apply this framework to public datasets together with newly generated PRO-seq and Micro-C data from primate immune cells, enabling a broad comparison of promoter-proximal pausing across conditions, cell types, and species. Analyses of these datasets revealed striking differences across transcriptional perturbations and distinct patterns of variation in pause-escape kinetics and pause-site distributions across cell types and species, with only weak coupling between them. Integration with chromatin and sequence features showed that lower pause-escape rates are associated with stronger promoter-proximal nucleosome occupancy, whereas changes in pause-site dispersion are associated with sequence features such as GC skew. Together, these results reveal kinetic and distributional dimensions of pausing variation across biological contexts.

## Introduction

The regulation of transcription is central to cellular identity, development, and evolution [1, 2]. Among the key regulatory checkpoints in metazoan transcription is promoter-proximal pausing of RNA polymerase II (Pol II), in which engaged polymerase accumulates approximately 20–60 nucleotides downstream of transcription start sites [3,4]. Promoter-proximal pausing functions as a rate-limiting step in early elongation and integrates developmental, environmental, and signaling inputs to fine-tune transcriptional output (reviewed in [5–7]). Understanding how promoter-proximal pausing varies across genes and biological contexts therefore remains an important goal in transcription research.

Nascent RNA sequencing technologies, including GRO-seq [8], PRO-seq [4, 9], and NET-seq [10–12], have enabled high-resolution measurements of transcriptionally engaged Pol II and revealed the genome-wide prevalence of promoter-proximal pausing [13, 14]. Much of the previous work on promoter-proximal pausing has focused on the extent of pausing, often quantified using descriptive measures such as the pausing index, defined as the ratio of promoter-proximal to gene-body polymerase signal [8, 15, 16]. These studies have established pause-escape as an important regulatory checkpoint in transcription. In particular, P-TEFb-mediated pause escape has emerged as a central mechanism controlling productive elongation, and perturbation of this pathway can profoundly affect transcriptional output and cellular responses [17–19].

In addition to variation in pausing kinetics, nascent RNA sequencing data reveal substantial variation in how paused polymerase is spatially distributed near promoters [4, 8, 20]. Thus, promoter-proximal pausing can be viewed as having distinct *kinetic* and *distributional* aspects: that is, aspects describing the rate at which polymerase escapes into productive elongation and aspects describing how paused polymerase is organized near promoters. Furthermore, perturbations such as NELF depletion can induce clear shifts in pause positions [21–23], suggesting that measures of pause release alone may only partially capture the relevant biology. The spatial organization of paused polymerase appears to represent an additional important dimension of promoter-proximal pausing.

Despite growing evidence that both kinetic and distributional variation contribute to promoter-proximal pausing, few methods are available for quantitatively characterizing these features across biological contexts. As a result, relatively little is known about how pause-site distributions vary across conditions, cell types, and species, or about how kinetic and distributional aspects of pausing are interrelated. These limitations, in turn, make it difficult to systematically relate variation in pausing to changes in chromatin organization, promoter architecture, and other regulatory features, impeding a full understanding of the mechanisms underlying this critical regulatory step.

To address these questions, we build on our previous probabilistic framework for estimating transcriptional dynamics from nascent RNA sequencing data [24, 25] and develop a series of likelihood-ratio tests (LRTs) for rigorous comparative analysis of pause-escape kinetics and pause-site distributions across biological contexts. Using simulated datasets, we evaluate the statistical performance of the proposed tests. We then apply this framework to perturbation datasets and newly generated Micro-C and nascent RNA sequencing data from primate immune cells to characterize variation in promoter-proximal pausing and to investigate its relationships with chromatin organization and promoter sequence composition. Altogether, we show that variation in promoter-proximal pausing can be described in terms of separate but interrelated kinetic and distributional axes, with different behaviors in different biological contexts.

## Results

### Likelihood-ratio tests detect changes in promoter-proximal pausing dynamics

We previously established a probabilistic framework for modeling transcriptional dynamics [24, 26]. Building on this foundation, we developed rigorous likelihood-ratio tests (LRTs) for differences in pause-escape rates (*β*) and pause-site distributions (*f_k_*) between biological conditions (**Fig. 1A**). These methods are implemented in STADyUM (Statistical Transcriptome Analysis under a Dynamic Unified Model), available through Bioconductor (see **Methods**). To our knowledge, these represent the first model-based tests specifically designed to detect changes in promoter-proximal pausing. Detecting such differences is critical for distinguishing whether changes in pausing reflect shifts in transcriptional kinetics, alterations in pause-site organization, or both.

**Figure 1:**
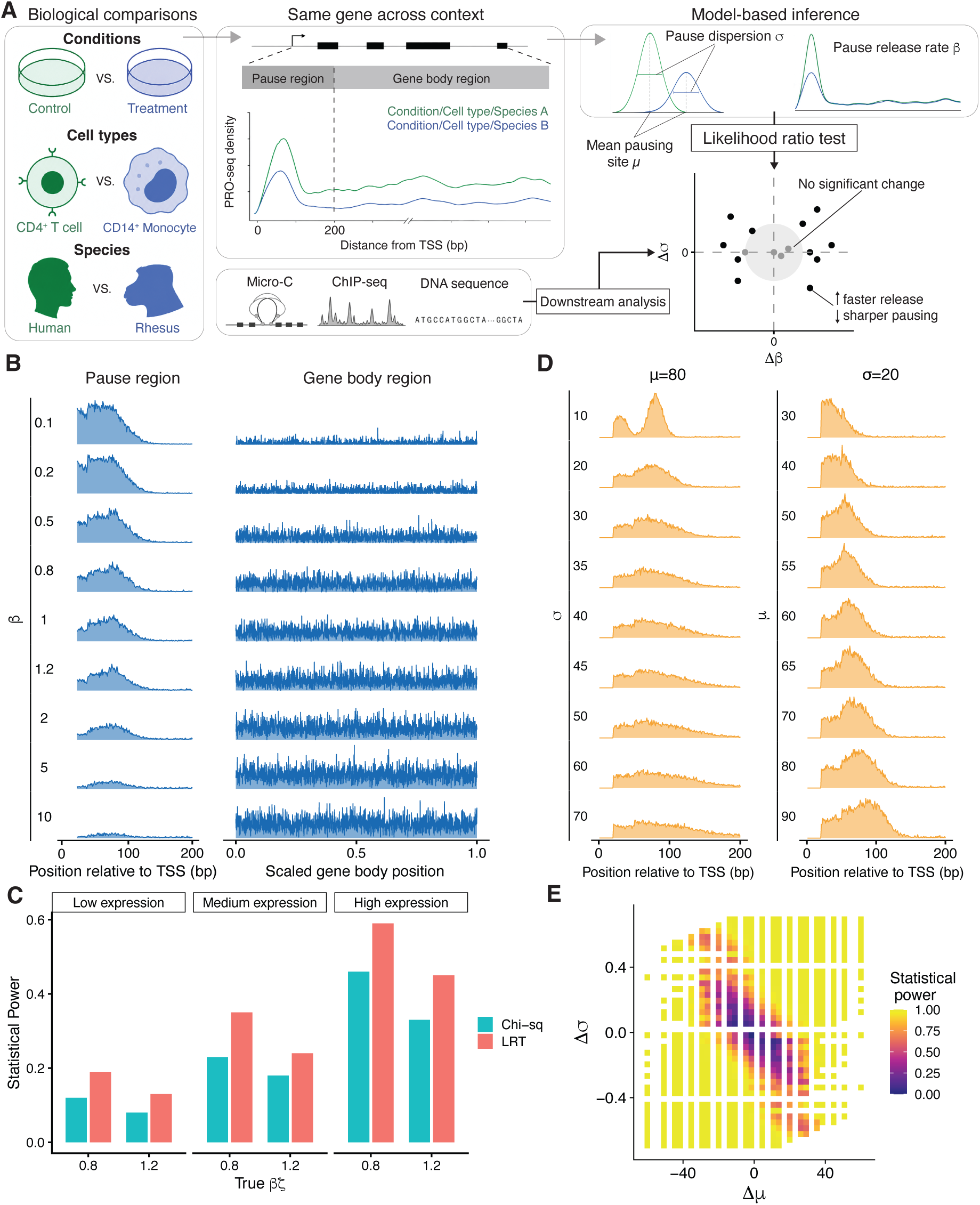
Likelihood-ratio testing framework for quantifying pause-escape kinetics and pausing distribution. **A.** Overview of the comparative probabilistic framework for promoter-proximal pausing analysis across conditions, cell types, and species. The framework uses likelihood-ratio tests to quantify changes in pause-escape kinetics and pausing distributions from nascent RNA sequencing data, followed by comparative analysis of chromatin and sequence features. **B.** Simulated PRO-seq profiles across different pause-escape rates (*β*), with fixed initiation rate and elongation rate (*ζ* = 2,000 nt per minute). **C.** Statistical power comparison between the LRT and chi-squared test for detecting changes in *β*. **D.** Representative simulated pause-site distributions with varying pause-site dispersion (*σ*, left) and mean position (*µ*, right). **E.** Statistical power surface of the distributional LRT across combinations of changes in *µ* and *σ*.

To evaluate the LRTs, we simulated nascent RNA profiles across a grid of parameter values ranging from 10-fold decreases to 10-fold increases relative to a reference setting. As expected, increasing *β* by 100-fold (from 0.1 to 10 events per minute), while holding initiation-rate fixed, caused simulated read counts to be increasingly redistributed from the promoter-proximal region into the gene body (**Fig. 1B**). For comparison, we benchmarked our likelihood-ratio tests against a simple chi-squared test based on relative pause and gene-body read densities, representing a minimal reference for detecting changes in pausing behavior [27]. Overall, we found that the LRT for *β* showed high sensitivity across simulated datasets and consistently outperformed the chi-squared test in detecting moderate (20%) changes in pause-escape rates (**Fig. 1C** & **Supplementary Fig. S1A–B**).

We next assessed the LRT for detecting differences in pause-site distributions by jointly varying the mean pause position (*µ*) and the dispersion of pause positions (*σ*; **Fig. 1D**). The test identified distributional changes across this two-parameter grid, yielding a two-dimensional power surface that was sensitive to coordinated shifts in both the location and spread of paused polymerase (**Fig. 1E** & **Supplementary Fig. S1C–D**). Together, these simulations demonstrate that our LRTs reliably detect changes in both pause-escape rates and pause-site distributions, providing a rigorous basis for comparative analyses of pausing dynamics.

### Distinct perturbations differentially affect pause-escape kinetics and pause-site distributions

Previous studies generally have not examined how different regulatory perturbations affect promoter-proximal pausing from both kinetic and distributional perspectives. We therefore reanalyzed previously published nascent transcription datasets following targeted disruption of pause-escape machinery [21]. We first applied our LRT to quantify changes in pause-escape rates following treatment with NVP-2, a widely used inhibitor of P-TEFb [21]. Genome-wide analysis revealed pervasive reductions in pause-escape rates after NVP-2 treatment, with more than 99% of genes showing significantly decreased *β* values (**Fig. 2A**), consistent with the established role of P-TEFb in promoting RNA Pol II pause escape [28]. In contrast, the overall spatial organization of promoter-proximal pausing remained largely preserved following treatment (**Fig. 2B**). When visualized jointly, NVP-2 treatment produced a highly directional shift dominated by changes in *β*, with relatively minor changes in pause-site dispersion (**Fig. 2C**). Similar patterns were observed following treatment with flavopiridol (**Supplementary Fig. S2A–C**), another well-established inhibitor of P-TEFb [29], suggesting that P-TEFb inhibition primarily affects pause-escape kinetics while having comparatively limited effects on pause-site distribution.

**Figure 2:**
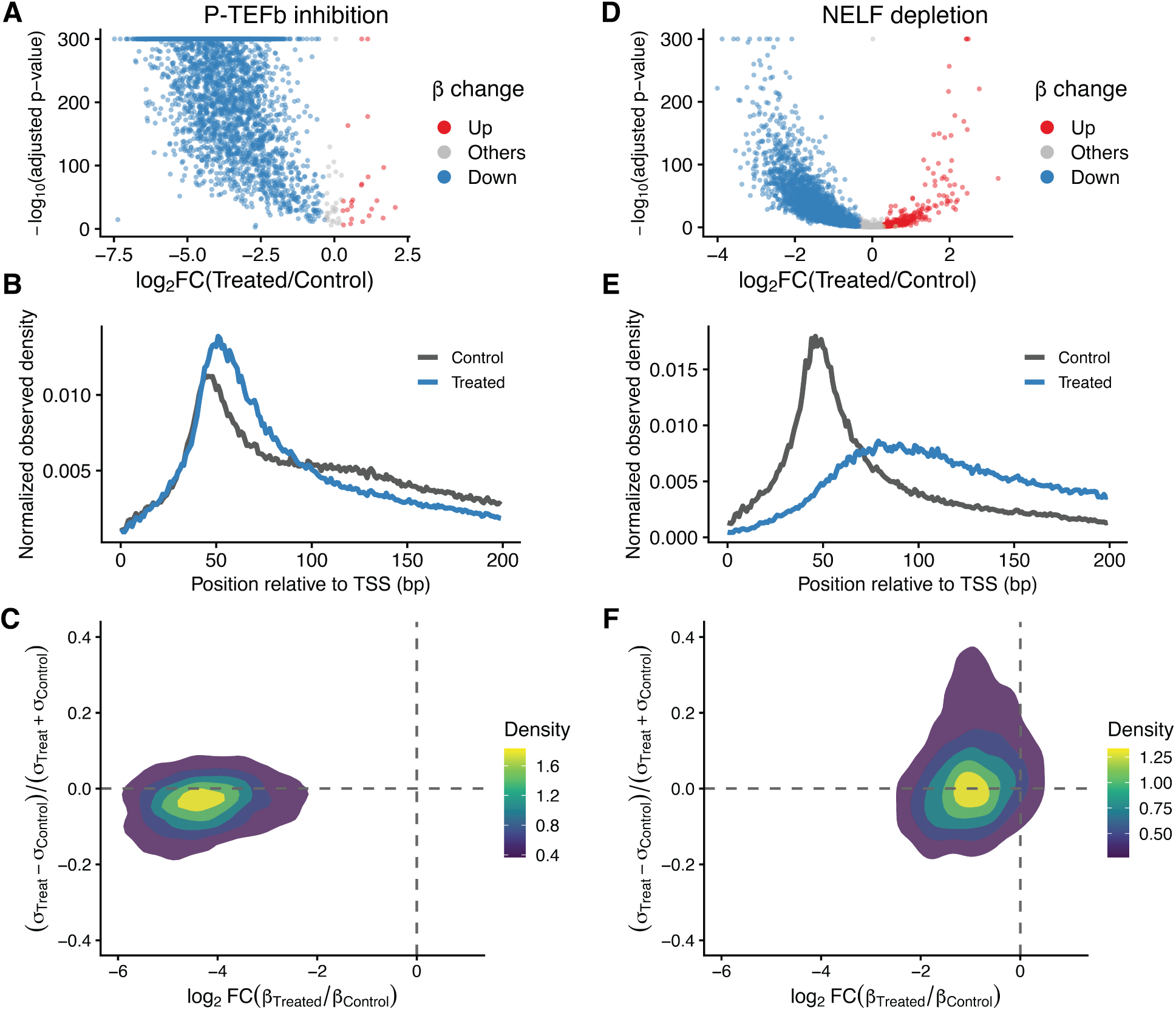
Perturbation experiments reveal distinct regulatory effects on pause-escape kinetics and pausing distribution. **A.** Genome-wide LRT results for pause-escape rates (*β*) following P-TEFb inhibition. **B.** Metaplots of normalized PRO-seq density downstream of TSS in control and NVP-2–treated cells. **C.** Global distributions of log_2_ fold changes in *β* and pausing dispersion *σ* under P-TEFb inhibition. **D.** Genome-wide LRT results for pause-escape rates (*β*) following NELF depletion. **E.** Metaplots of normalized PRO-seq density downstream of TSS in control and NELF-depleted cells. **F.** Global distributions of log_2_ fold changes in *β* and pausing dispersion *σ* under NELF depletion.

We next examined the effects of acute depletion of NELF, a core component of the promoter-proximal pausing machinery [21]. Genome-wide LRT analysis revealed widespread changes in *β* following NELF depletion, with 75.6% of genes showing significantly decreased *β* values and 9.2% showing significantly increased *β* values (**Fig. 2D**). Metaplot analysis further showed that NELF depletion shifted paused Pol II distributions downstream and broadened promoter-proximal pausing patterns (**Fig. 2E**), consistent with previous observations [21, 22]. Joint analysis of *β* and *σ* showed that NELF depletion produced substantial changes in both pause-escape kinetics and pause-site dispersion (**Fig. 2F**). Notably, genes exhibiting broader pause-site distributions frequently also showed reduced pause-escape rates, suggesting partial coupling between changes in pausing dispersion and pause-escape kinetics following NELF depletion. Taken together, joint analysis of pause-escape kinetics and pause-site distribution suggests that different transcriptional perturbations preferentially affect distinct aspects of promoter-proximal pausing: P-TEFb inhibition primarily reduces pause-escape rates, whereas NELF depletion extensively reorganizes pause-site distributions while exerting mixed effects on *β*.

### Distinct chromatin features are associated with different aspects of promoter-proximal pausing within cell types

Having shown that pause-escape kinetics (*β*) and pause-site dispersion (*σ*) respond differentially to regulatory perturbation, we next asked how these two aspects of pausing vary across cell types within species and across species within matching cell types. To address this question, we generated matched PRO-seq and Micro-C datasets in humans (*Homo sapiens*), rhesus macaques (*Macaca mulatta*), and olive baboons (*Papio anubis*) for CD4^+^ T cells and CD14^+^ monocytes (see **Methods**). In this section, we examine human CD4^+^ T cells as a representative system to characterize the relationships between pause-escape kinetics, pause-site distribution, and promoter chromatin organization.

Within human CD4^+^ T cells, the pause-escape rate (*β*) positively co-varies with transcriptional activity (*χ*, defined as the average read density in the gene body; Spearman’s *ρ* = 0.61, *N* = 7, 029; see **Methods**), consistent with its estimation from ratios of gene-body to pause-peak signals [24] (**Fig. 3A**). Similar relationships were observed across species and cell types, with Spearman’s *ρ* values ranging from 0.60 to 0.71 (**Supplementary Fig. S3A**). For subsequent analyses, genes were grouped by *β* into quintiles (Q1–Q5, from lowest to highest) and by *χ* into tertiles (Low, Medium, and High).

**Figure 3:**
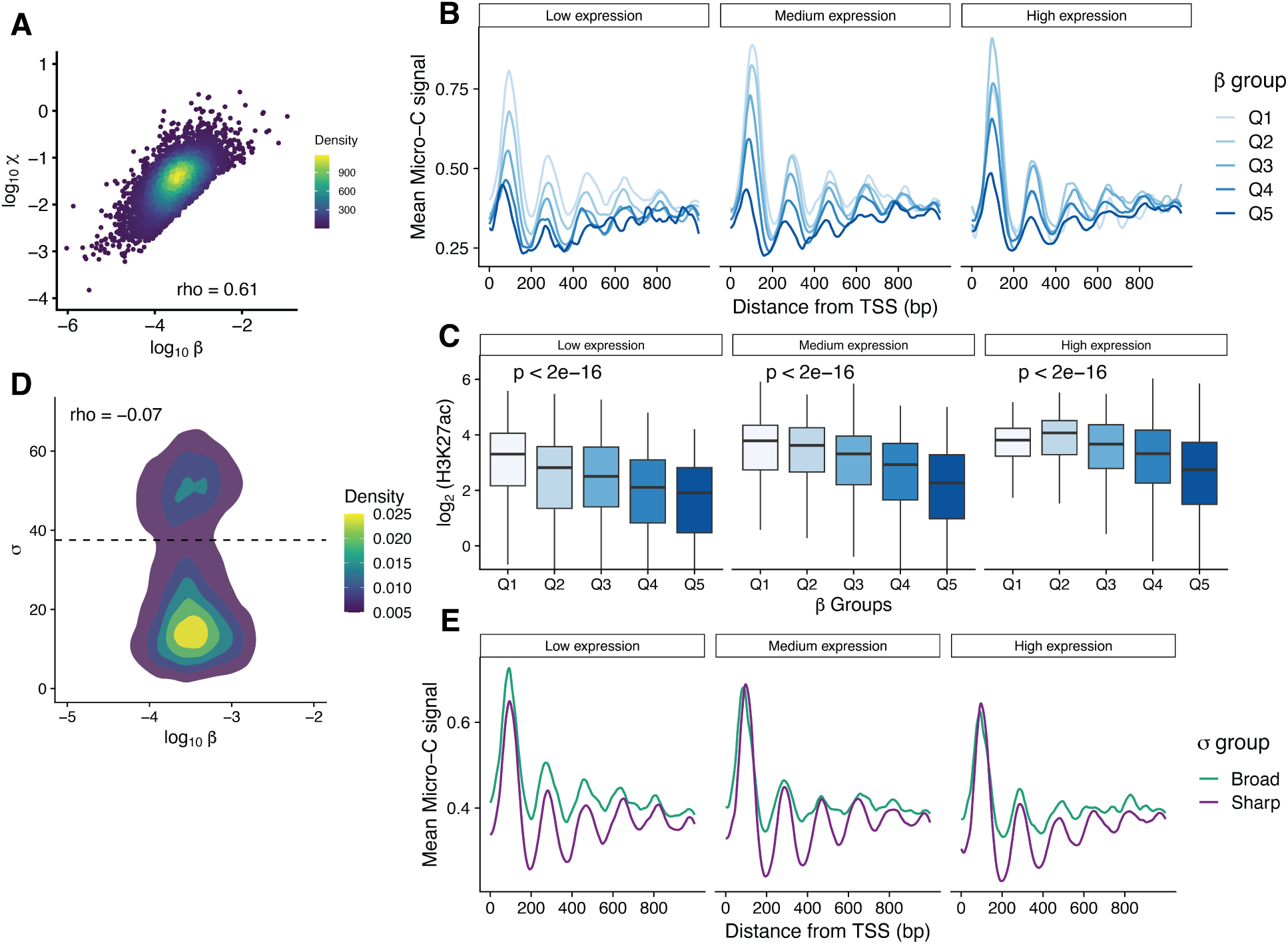
Promoter-proximal pausing can be described using separable kinetic and distributional aspects. **A.** Relationship between pause-escape rate (*β*) and transcriptional activity (*χ*) across genes in human CD4^+^ T cells. The Spearman correlation coefficient is indicated. Three extreme outliers with unusually large *β* values are omitted for visualization clarity. **B.** Average Micro-C signal around TSSs for genes stratified by *β* quintiles (Q1–Q5) in CD4^+^ T cells. Genes are further grouped by transcription activity *χ* (low, medium, and high). **C.** Distributions of log_2_ H3K27ac signal at promoters for genes stratified by *β* quintiles (Q1–Q5) and transcriptional activity groups (*χ*). *P* values were calculated using Kruskal–Wallis tests across the five *β* groups. **D.** Relationship between pause-escape rate (*β*) and pausing dispersion (*σ*) in human CD4^+^ T cells. The dashed line indicates the threshold separating sharp and broad pausing dispersion. **E.** Average Micro-C signal around TSSs for genes grouped by pausing dispersion (*σ*; Broad versus Sharp) across transcriptional activity groups.

Promoter-proximal nucleosome organization has been implicated in shaping early elongation dynamics [30–32]. To examine the relationship between pause-escape kinetics and promoter-proximal nucleosome organization, we extracted one-dimensional Micro-C signals across the 1 kb region downstream of the TSS (see **Methods**). Under the *χ* stratification, genes with lower *β* exhibit stronger promoter-proximal nucleosome occupancy and clearer positioning patterns in human CD4^+^ T cells (**Fig. 3B**). Similar relationships were also observed across species and cell types (**Supplementary Fig. S3B**).

Previous studies have shown that different promoter motifs (such as TATA box, GAGA factor, and CpG islands) are associated with distinct promoter-proximal nucleosome organization patterns [33–35], suggesting that promoter sequence context could potentially confound the relationship between *β* and nucleosome occupancy. Consistent with previous observations, nucleosome organization downstream of the TSS varies across promoter classes (**Supplementary Fig. S4A–B**). More specifically, genes containing only the CCAAT motif exhibit the strongest +1 nucleosome occupancy among the promoter classes examined. However, after stratifying genes by promoter class, lower *β* remains consistently associated with stronger nucleosome occupancy across promoter categories (**Supplementary Fig. S4C**; see **Discussion**). Similar patterns were also observed in human CD14^+^ monocytes (**Supplementary Fig. S4D–F**).

Because active promoter-associated histone marks such as H3K27ac and H3K4me3 are enriched on promoter-proximal nucleosomes [36], we next examined their relationships with pause-escape kinetics. Using ENCODE ChIP-seq datasets [37], we extracted histone modification signals across the ±500 bp region surrounding the TSS (see **Methods**). Under the *χ* stratification, genes with lower *β* exhibit stronger H3K27ac and H3K4me3 signals (**Fig. 3C** & **Supplementary Fig. S4G–I**), consistent with the associations between *β* and nucleosome observed in **Fig. 3B**.

We next investigated whether pausing dispersion (*σ*) exhibits distinct relationships with transcriptional activity and promoter chromatin organization. *σ* shows little correlation with pause-escape rate (*β*) (**Fig. 3D** and **Supplementary Fig. S5A**), suggesting that pause-site distribution captures a distinct aspect of promoter-proximal pausing. In contrast, *σ* and *µ* remain strongly correlated within immune cells (**Supplementary Fig. S5B**). To examine the relationship between pausing dispersion and promoter chromatin organization, we separated genes into sharp and broad pausing groups (**Fig. 3D** & **Supplementary Fig. S6A**, see **Methods** for details of *σ* grouping). Genes with sharp pause peaks exhibit clearer and more regularly positioned promoter-proximal nucleosomes, whereas broad pausing is associated with less well-defined positioning patterns (**Fig. 3E** and **Supplementary Fig. S6B**). A similar pattern is observed after stratifying genes by *β*, indicating that the relationship between pausing dispersion and nucleosome organization is not restricted to genes with high or low pause-escape rates (**Supplementary Fig. S6C**). These observations suggest that variation in *σ* is associated with differences in promoter-proximal organization rather than simply tracking *β* or transcriptional activity.

Together, these results show that, within cell types, pause-escape kinetics and pause-site distributions exhibit distinct relationships with promoter chromatin organization. Genes with lower pause-escape rates tend to exhibit stronger promoter-proximal nucleosome occupancy and higher levels of active promoter-associated histone marks across promoter classes and transcriptional activity levels, whereas sharp pausing distributions are associated with more phased and better positioned +1 nucleosomes, largely independent of transcriptional activity and pause-escape rate. These within-cell-type associations provide biological context for interpreting *β* and *σ* as distinct aspects of promoter-proximal pausing.

### Cell-type comparisons reveal distinct patterns of variation in pause-escape kinetics and pause-site distribution

To determine how promoter-proximal pausing parameters vary across cellular contexts, we compared parameter estimates between human CD4^+^ T cells and CD14^+^ monocytes. Rank-based comparison of *β* quintiles revealed strong enrichment along the diagonal of the CD4–CD14 matrix (**Supplementary Fig. S7A**), with similar patterns also observed in rhesus macaques and olive baboons (**Supplementary Fig. S7B–C**), indicating substantial preservation of global kinetic ordering across cell types. Functional enrichment analysis further showed that genes within individual *β* strata are associated with coherent biological processes (**Supplementary Fig. S7D–E**). Genes with small *β* are enriched for metabolic processes, whereas genes with large *β* are associated with signaling pathways involved in immune cell activation, consistent with previous observations linking stronger promoter-proximal pausing to metabolic and housekeeping functions [15]. Despite this global conservation, our LRT analysis identified widespread gene-level changes in *β* between human CD4^+^ T cells and CD14^+^ monocytes, with 25.8% of genes showing significantly increased *β* values and 36.3% showing significantly decreased *β* values (**Fig. 4A**). Similar patterns were also observed in rhesus macaques and olive baboons, where 63.8% and 82.1% of genes exhibited significant changes in *β* between cell types, respectively (**Supplementary Fig. S7F–G**). Together, these results suggest that pause-escape kinetics maintain a globally stable organization while retaining substantial regulatory plasticity across cellular contexts.

**Figure 4:**
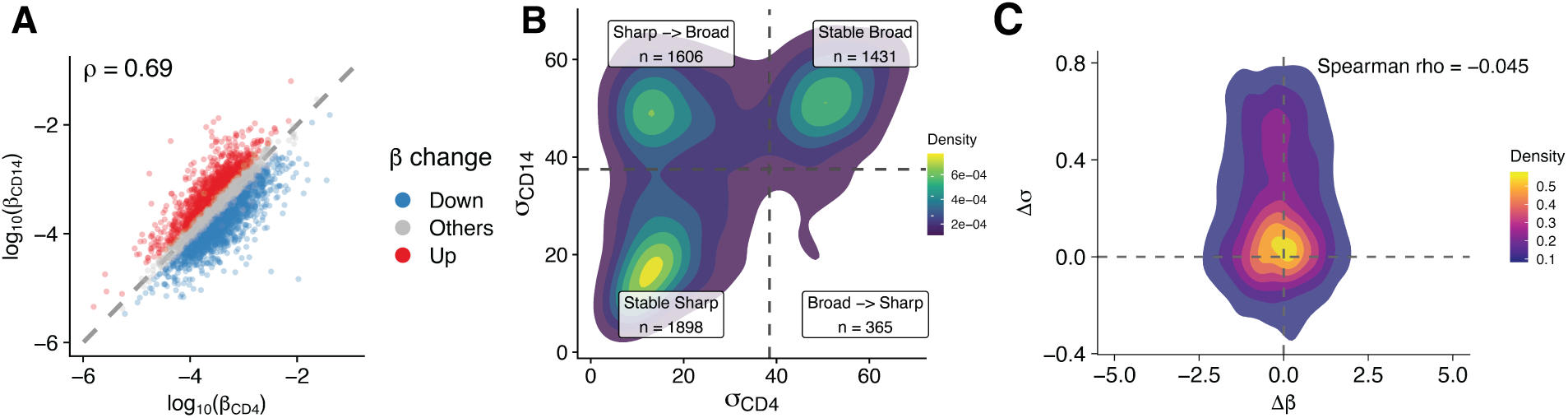
Cell-type differences in pause-escape rate and pausing distribution. **A.** Comparison of pause-escape rates (*β*) between human CD4^+^ T cells and CD14^+^ monocytes. Genes are colored according to LRT-inferred significant changes in *β*. **B.** Comparison of pausing dispersion (*σ*) between human CD4^+^ T cells and CD14^+^ monocytes. Genes are classified according to broad and sharp pausing dispersion (*σ*) in each cell type. **C.** Joint distribution of changes in pause-escape rate (Δ*β*) and pausing dispersion (Δ*σ*) across cell types. The Spearman correlation coefficient is shown.

We next examined pausing dispersion (*σ*) between human CD4^+^ T cells and CD14^+^ monocytes. Although *σ* values show global alignment between the two cell types, substantial gene-level deviations from the diagonal are evident (**Fig. 4B**). Stratifying genes by sharp and broad pausing groups in each cell type reveals four classes of change. Notably, approximately 30.3% (1606 out of 5300) of genes switch between sharp and broad pausing groups between the two cell types, indicating that variation in pausing dispersion frequently involves changes between sharper and broader pause-site distributions. Similar sharp–broad changes were consistently observed in rhesus macaques and olive baboons (**Supplementary Fig. S8A–B**), suggesting that this pattern of pausing-dispersion variation is shared across primate species. To compare kinetic and distributional changes directly, we mapped shifts in *β* against shifts in *σ*. Although individual genes display changes in both parameters, their joint distribution shows minimal overall association across humans, rhesus macaques, and olive baboons (**Fig. 4C** and **Supplementary Fig. S8C–D**), indicating weak coupling between kinetic and distributional changes.

Together, these analyses suggest that pause-escape kinetics and pause-site distribution exhibit distinct modes of variation across cellular contexts. While the relative ordering of genes by pause-escape rate remains largely preserved across cell types, pause-escape kinetics retain substantial regulatory plasticity. Variation in pausing dispersion more frequently involves changes between sharper and broader pause-site distributions. The weak association between changes in *β* and *σ* further supports the view that kinetic and distributional aspects of pausing can vary differently across cellular contexts.

### Evolutionary comparisons reveal conservation of pause-escape kinetics and structured variation in pause-site distribution

To extend this comparative framework across evolutionary timescales, we compared parameter estimates between human and rhesus CD4^+^ T cells using one-to-one orthologous genes. Orthologous loci were stratified into conserved and non-conserved TSS groups based on cross-species alignment of promoter-proximal pausing regions (**Fig. 5A**, see also **Methods**). This classification defines evolutionary preservation of promoter-proximal architecture independently of parameter magnitude.

**Figure 5:**
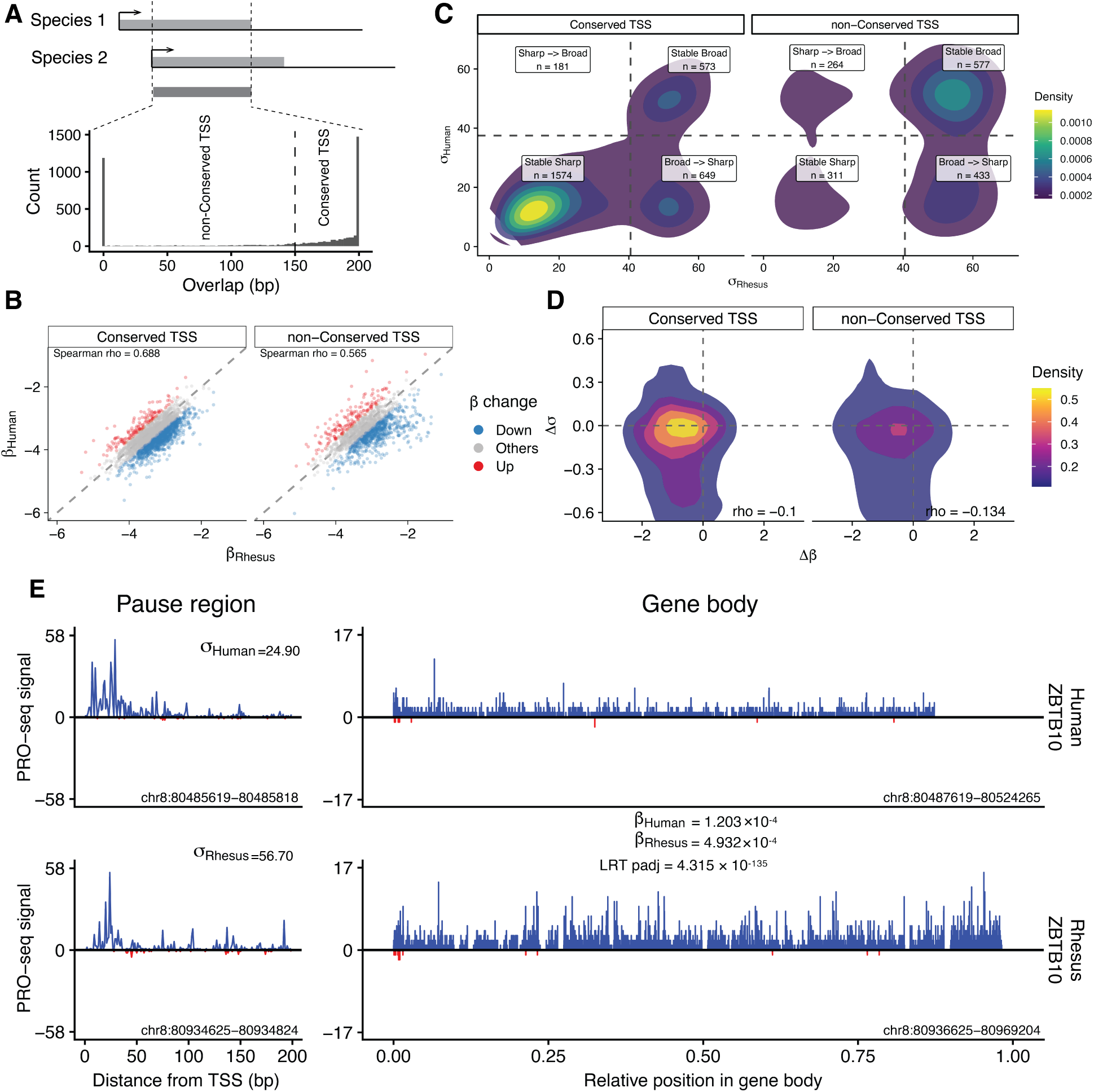
Cross-species changes in pause-escape rate and pausing distribution. **A.** Classification of orthologous genes into conserved and non-conserved transcription start sites (TSSs) based on overlap of promoter-proximal pausing regions across species. **B.** Comparison of pause-escape rates (*β*) between orthologous genes in human and rhesus CD4^+^ T cells, colored by LRT-inferred changes in *β*. **C.** Comparison of pausing dispersion (*σ*) between orthologous genes across species. Genes are classified according to broad and sharp pausing dispersion (*σ*) in each species. **D.** Joint distribution of changes in pause-escape rate (Δ*β*) and pausing dispersion (Δ*σ*) across species. The Spearman correlation coefficients are shown. **E.** Representative gene illustrating cross-species differences in pause-escape kinetics and pausing distributions.

Across orthologous genes, *β* exhibits a positive global correlation between species (**Fig. 5B**), indicating that not only overall transcriptional output but also the relative balance between promoter-proximal pausing and productive elongation is similar across species. This pattern suggests that pause-escape kinetics show reproducible cross-species alignment beyond expression level alone. Although the overall ordering of *β* values remains globally preserved across species, our LRT analysis identified widespread gene-level differences in pause-escape kinetics. Notably, 39.8% and 42.9% of genes exhibited significantly decreased *β* values among conserved TSS and non-conserved TSS genes, respectively. In addition, conserved TSS genes retained stronger cross-species correlation, whereas non-conserved loci exhibited greater dispersion in *β*. Similar patterns were consistently observed across other cross-species comparisons (**Supplementary Fig. S9A**).

A parallel comparison of pausing dispersion (*σ*) reveals similar global persistence of pause-site organization, with 66.4% of genes retaining the same pausing-dispersion category across species (**Fig. 5C**), indicating substantial conservation of pause-site distribution. However, stratification by TSS conservation status revealed markedly greater changes between pausing-dispersion groups among non-conserved TSSs relative to conserved TSSs (45.0% versus 27.5%), suggesting that evolutionary remodeling of promoter architecture is associated with increased plasticity in pause-site distribution. Similar patterns were consistently observed across other cross-species comparisons (**Supplementary Fig. S9B**). To jointly visualize cross-species variation in pause-escape kinetics and pause-site distribution, we mapped shifts in *β* against shifts in *σ* (**Fig. 5D**). As expected, genes with conserved TSSs cluster near the origin, indicating relatively small changes in both parameters, whereas non-conserved loci exhibit substantially broader dispersion in both parameters. Notably, changes in *β* and *σ* show weak Spearman correlation across genes, a pattern consistently observed across other cross-species comparisons (**Supplementary Fig. S9C**). Representative loci illustrate how these shifts in *β* and *σ* appear at individual genes (**Fig. 5E**).

Together, these analyses indicate that although the overall organization of pause-escape kinetics and pause-site distribution remains broadly conserved across species, non-conserved TSSs exhibit substantially greater divergence in both pausing parameters, suggesting that evolutionary remodeling of promoter architecture is associated with increased plasticity in promoter-proximal pausing dynamics. Moreover, the weak correlation between cross-species changes in *β* and *σ* indicates that pause-escape kinetics and pause-site distribution show different patterns of cross-species variation.

### Chromatin features are associated with pause-escape kinetics and pause-site distribution

Having observed distinct patterns of variation in pause-escape kinetics and pause-site distribution across cell types and species, we next asked which promoter features are associated with these two aspects of promoter-proximal pausing. We first investigated how changes in pause-escape kinetics relate to promoter-proximal chromatin organization, focusing on the +1 nucleosome located within the first 200 bp downstream of the TSS (see **Methods**). Because promoter-proximal nucleosome occupancy is strongly associated with transcriptional activity (**Supplementary Fig. S10A**), genes were stratified by transcriptional activity (*χ*). Across all expression strata, reduced +1 nucleosome occupancy is consistently associated with increased pause-escape rates (*β*) (**Fig. 6A**; see also refs. [21,31,32,38]), a relationship also observed in rhesus macaque (**Supplementary Fig. S10B–C**).

**Figure 6:**
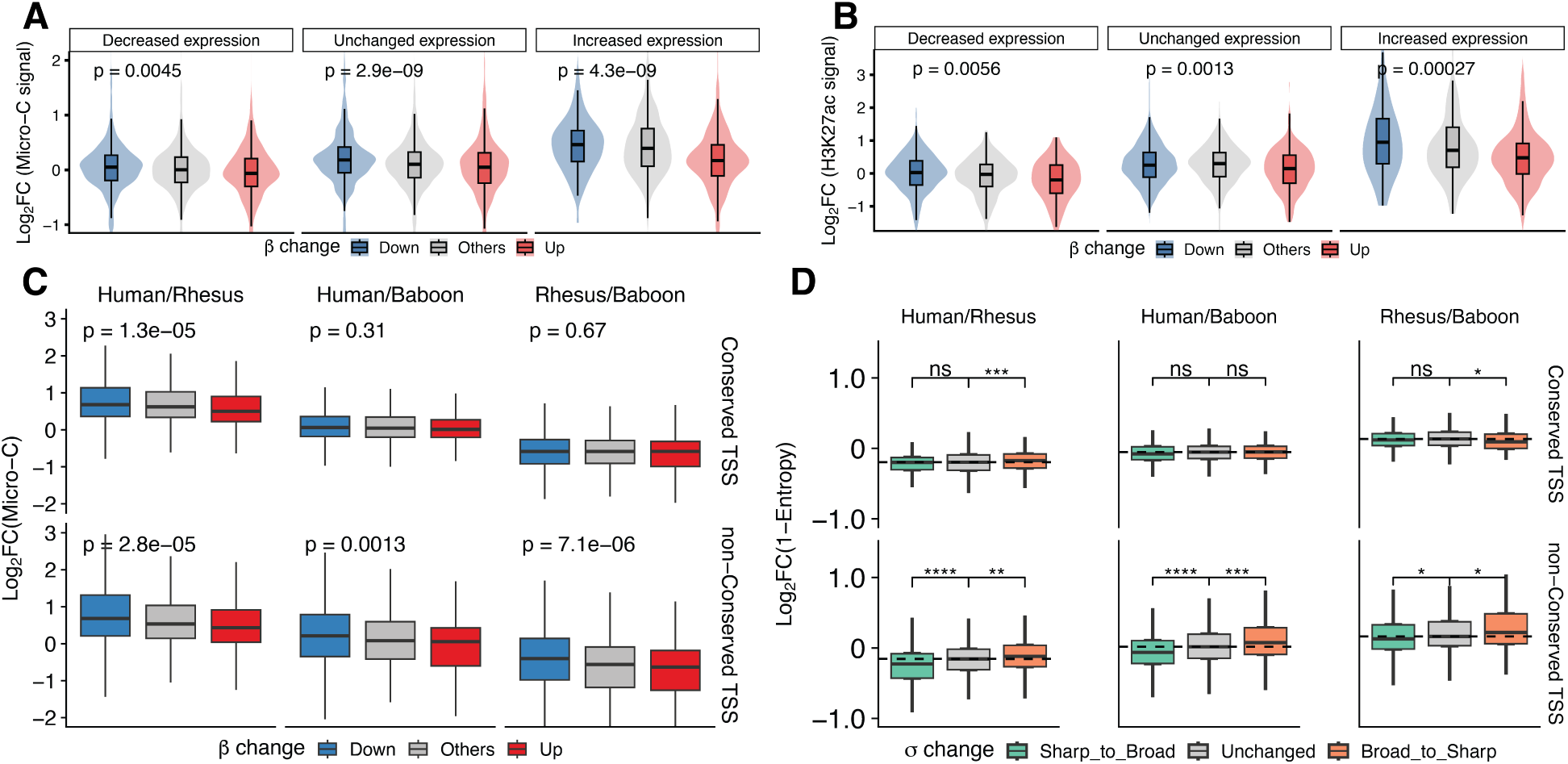
Chromatin features are associated with variation in pause-escape kinetics and pausing distributions. **A.** Changes in +1 nucleosome occupancy across genes with different changes in pause-escape rate (*β*) across cell types. Genes are grouped by transcriptional activity change. **B.** Changes in H3K27ac signal across genes with different changes in pause-escape rate (*β*). Genes are grouped as panel A. **C.** Cross-species changes in +1 nucleosome occupancy associated with changes in pause-escape rate (*β*). Orthologous genes are stratified by TSS conservation status. In each comparison (e.g., Human/Rhesus), log fold changes are calculated with the first species as the numerator and the second as the denominator. **D.** Changes in nucleosome positioning entropy associated with changes in pausing dispersion (*σ*) across species. For panels A–C, P values were calculated using Kruskal–Wallis tests across the three *β*-change groups. For panel D, significance was assessed using pairwise Wilcoxon tests with adjusted *P* values. ns, not significant; * adjusted *P <* 0.05; ** adjusted *P <* 0.01; *** adjusted *P <* 0.001; **** adjusted *P <* 0.0001.

We next examined promoter-associated histone modifications. Although local changes in both H3K4me3 and H3K27ac are positively associated with transcriptional activity (**Supplementary Fig. S10D-E**), only H3K27ac shows a consistent negative association with *β* after stratification by *χ* (**Fig. 6B**), whereas H3K4-me3 does not (**Supplementary Fig. S10F**). Together, these results identify +1 nucleosome occupancy and H3K27ac as promoter-proximal chromatin features particularly associated with variation in pause-escape kinetics.

We next asked whether pause-escape kinetics and promoter-proximal nucleosome occupancy exhibit coordinated evolutionary changes across species. Cross-species changes in +1 nucleosome occupancy show little association with changes in *χ* (**Supplementary Fig. S10G**), and therefore expression stratification was not applied in subsequent analyses. Orthologous genes were stratified into conserved and non-conserved TSS groups, as in **Fig. 5A**. Among conserved TSS genes, decreases in +1 nucleosome occupancy are accompanied by increases in *β* only in the human–rhesus comparison in CD4^+^ T cells, whereas other species comparisons show no consistent directional trend (**Fig. 6C**). In contrast, among non-conserved TSS genes, decreases in +1 nucleosome occupancy are consistently associated with increases in *β* across comparisons. These results suggest that evolutionary changes in pause-escape kinetics are more tightly coupled to changes in promoter-proximal nucleosome occupancy at non-conserved TSS genes.

We also examined how pausing dispersion relates to nucleosome positioning architecture. Nucleosome positioning within the 0–200 bp region downstream of the TSS was quantified using an entropy-based metric (see **Methods**). Paired comparisons across cell types did not reveal statistically significant differences in +1 nucleosome positioning across changes between sharp and broad pausing groups (**Supplementary Fig. S10H**). We next performed the same analysis for cross-species comparisons. Among genes with conserved TSS, changes in pausing distribution show little association with nucleosome positioning differences across species (**Fig. 6D**). In contrast, among genes with non-conserved TSS, changes from sharp to broad pausing tend to coincide with weaker +1 nucleosome positioning, whereas broad-to-sharp changes show the opposite pattern.

Together, these analyses reveal distinct chromatin associations for the kinetic and distributional aspects of promoter-proximal pausing. Pause-escape kinetics show consistent relationships with promoter-proximal chromatin features. In particular, +1 nucleosome occupancy is associated with variation in *β* across both cellular and evolutionary comparisons, whereas H3K27ac shows a similar relationship across cellular contexts. In contrast, associations involving pausing dispersion are weaker and become most apparent at genes with non-conserved TSS, where changes in pausing dispersion are accompanied by altered +1 nucleosome positioning.

### Sequence features are associated with pause-escape kinetics and pause-site distribution

Previous studies have suggested that GC-rich promoter regions and related sequence features can influence Pol II pausing through effects on DNA stability and secondary structure formation [39–41]. We therefore examined whether sequence features near the TSS, particularly GC content and GC skew, are associated with variation in pause-escape kinetics and pause-site distribution across species.

We first examined the relationship between pause-escape kinetics and GC content in the promoter-proximal pausing region. GC content was calculated within the 1000 bp region surrounding the TSS for each active gene and then averaged across genes within each species-cell-type combination to capture promoter-proximal sequence composition (**Fig. 7A**). Consistent with previous observations, pausing regions showed high GC content across all cell types [42], with baboon genes showing slightly lower GC content than the other species. As expected, GC content shows little change among genes with conserved TSS even when *β* differs significantly across species (**Fig. 7B**), indicating that kinetic divergence at these loci is largely decoupled from local sequence composition. In contrast, among non-conserved TSS genes, those exhibiting relatively lower *β* in human and rhesus macaque also showed markedly higher GC content within pausing regions compared with their baboon orthologs, a pattern observed in both CD14^+^ monocytes and CD4^+^ T cells (**Fig. 7B**). These results suggest that evolutionary changes in GC content may contribute to cross-species variation in *β*, potentially through effects on DNA stability and local nucleic acid structure that influence promoter-proximal pause escape.

**Figure 7:**
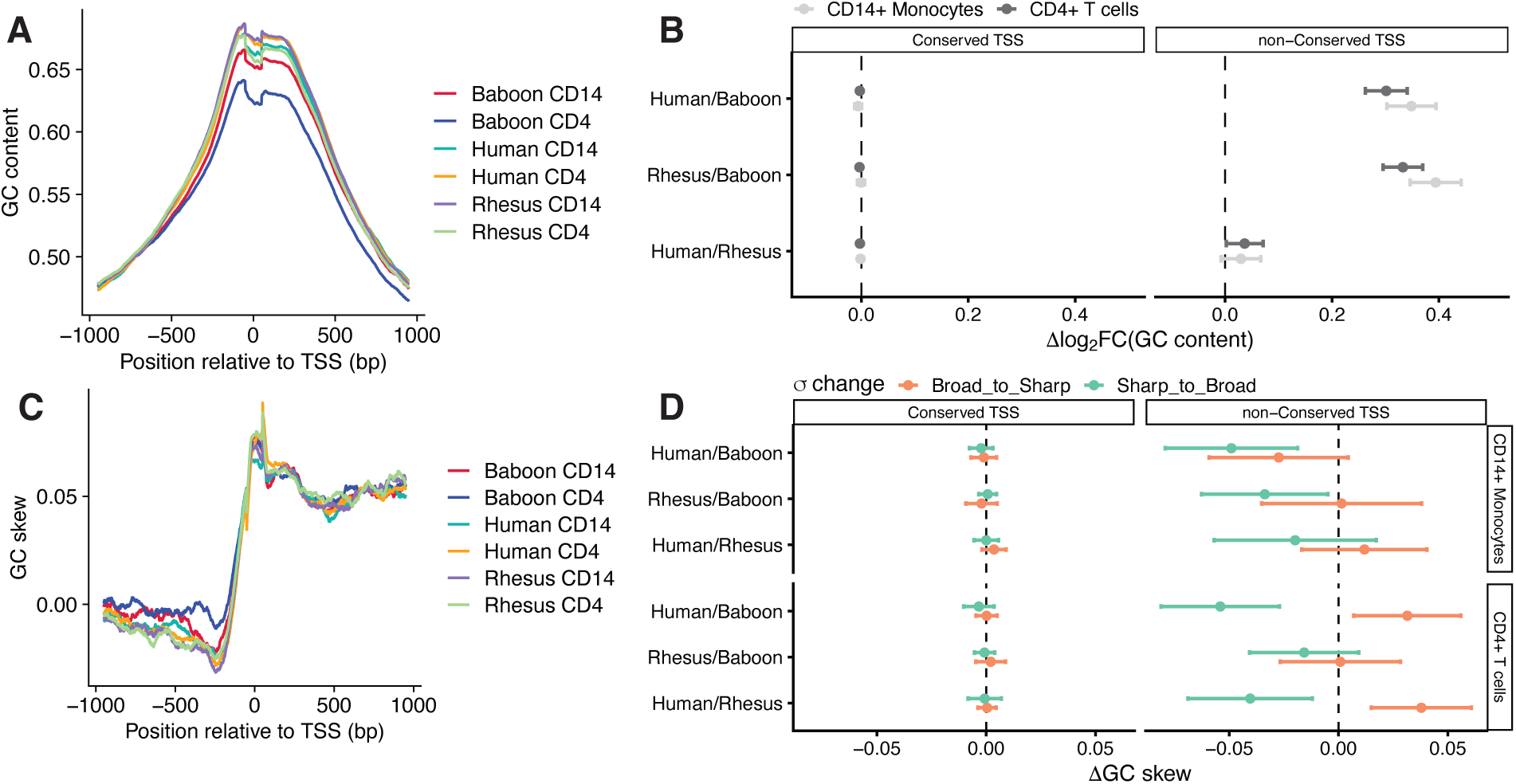
Promoter sequence features are associated with variation in pause-escape kinetics and pausing distributions. **A.** Average promoter-proximal GC content of active genes across species and cell types. **B.** Changes in GC content associated with changes in pause-escape rate (*β*) across species and TSS conservation groups. Effect sizes are defined as the difference in mean log_2_ fold change (GC content) between *β*_Down_ genes and genes with stable *β* (*β*_Others_) across species and TSS conservation groups. **C.** Average promoter-proximal GC skew of active genes across species and cell types. **D.** Changes in GC skew associated with changes in pausing dispersion (*σ*). Effect sizes are defined as the difference in mean GC skew change between genes with stable and changing pausing dispersion. For panels A and C, values at each position represent the mean GC content or GC skew within a 100-bp window (±50 bp) across all active genes in the corresponding species and cell type.

When stratifying genes by cross-species changes in pausing dispersion (*σ*), conserved TSS genes again show little variation in GC content across species (**Supplementary Fig. S11A**). In contrast, among non-conserved TSS genes, a clear association emerges between *σ* and GC content in CD4^+^ T cells. This pattern is more pronounced in human–baboon and human–rhesus comparisons and less evident in baboon–rhesus comparisons. Genes that change from broad to sharp pausing distributions tend to show increased GC content across species, whereas genes that change from sharp to broad show decreased GC content.

Because pausing distribution reflects the spatial locations of paused polymerases, we next examined whether sequence asymmetry near the TSS is associated with variation in pausing distribution. GC skew was calculated across the promoter-proximal region to quantify strand asymmetry in nucleotide composition, a feature of promoter architecture that has been linked to transcription initiation and R-loop-prone sequence environments [43]. Consistent with previous observations [42, 44], pausing regions showed high GC skew across all cell types (**Fig. 7C**). GC skew does not show a clear association with cross-species differences in *β* (**Supplementary Fig. S11B**). However, when stratifying genes by changes in *σ*, GC skew follows a directional trend similar to that observed for GC content in CD4^+^ T cells: sharp-to-broad changes are accompanied by reduced GC skew, whereas broad-to-sharp changes show increased GC skew (**Fig. 7D**). This pattern is less evident in CD14^+^ monocytes. These observations suggest that promoter-proximal GC skew is associated with variation in pausing distribution, potentially through effects on R-loop formation.

Together, these results indicate that cross-species variation in promoter-proximal pausing is accompanied by changes in local sequence composition. GC content is more strongly associated with variation in pause-escape kinetics (*β*), whereas GC skew shows a clearer relationship with pausing dispersion (*σ*), particularly at evolutionarily divergent promoters.

## Discussion

Promoter-proximal pausing has long been recognized as a key regulatory step in early transcription elongation and has been implicated in diverse biological processes, including transcriptional control, enhancer and promoter communication, viral infections, and developmental transitions [5, 6, 19, 45, 46]. Despite extensive mechanistic investigation, pausing behavior has typically been examined at the levels of individual genes or specific perturbations, without the benefit of quantitative modeling of its global organization across genes and biological contexts [24, 25]. Here we developed a comparative probabilistic framework that jointly models pause-escape kinetics and pause-site distributions, enabling systematic comparisons of promoter-proximal pausing across perturbations, cell types, and species. In addition, the framework also enables analysis of how kinetic and distributional properties change together across biological contexts, thereby providing a broader view of pausing dynamics than previously possible. To facilitate broader application, these methods are implemented in the STADyUM R package.

Application of the framework to acute NELF depletion and P-TEFb inhibition datasets further illustrates how comparative modeling of pausing dynamics can provide a more detailed view of transcriptional perturbation responses. Acute NELF depletion produced coordinated changes in both pause-site distributions and pause-escape kinetics, whereas P-TEFb inhibition primarily affected pause-escape kinetics, consistent with their established roles in stabilizing paused polymerases and promoting pause-escape, respectively [21, 28]. These observations suggest that the framework can help to reveal how different perturbations affect kinetic and distributional aspects of promoter-proximal pausing. Given that promoter-proximal pausing is linked to diverse regulatory processes involving chromatin remodeling [47], histone modification [48, 49], elongation control [50], and RNA-processing-associated factors [51], this type of comparative analysis may help evaluate whether perturbations of different regulatory factors preferentially affect pause-escape kinetics, pause-site distributions, or both.

Comparative analyses across the datasets generated in this study further revealed that pause-escape kinetics and pause-site distribution exhibit distinct modes of variation across biological contexts. Although the relative ordering of genes by pause-escape rate remained broadly preserved across cell types and species, widespread gene-level changes indicate substantial regulatory plasticity in pause-escape kinetics. By contrast, variation in pausing dispersion more frequently involved changes between sharper and broader pause-site distributions. These observations suggest that kinetic and distributional aspects of promoter-proximal pausing capture different patterns of variation. Notably, changes in *β* and *σ* were often only weakly coupled across cell-type and species comparisons, suggesting that pause-escape kinetics and pause-site distribution may be shaped by partially distinct regulatory and evolutionary influences.

Across within-cell-type, between-cell-type, and cross-species comparisons, lower pause-escape rates were generally associated with increased +1 nucleosome occupancy. These observations suggest that promo-ter-proximal chromatin organization is linked to pause-escape kinetics. Previous models have often viewed the +1 nucleosome as a physical barrier to early transcription elongation [32, 52, 53]. Our analyses are consistent with a view in which promoter-proximal nucleosome organization is associated with pause-escape kinetics, although their causal relationship remains unclear. In addition, decreased active promoter-associated histone modifications such as H3K27ac were also associated with increased pause-escape rates, although the mechanistic roles of active histone marks in promoter-proximal pausing remain incompletely understood [48, 49, 54–56].

Pausing dispersion reflects the spatial organization of paused polymerase near promoters. In the analyses presented here, sharp pausing distributions generally co-occurred with more phased and better-positioned +1 nucleosomes, whereas broader pausing distributions were more often associated with less constrained nucleosome organization. These observations link pausing dispersion to promoter-proximal nucleosome organization, but whether and how nucleosome positioning influences pause-site distribution remains an open question. In addition, how variation in pause-site distribution relates to downstream regulatory processes will be an important direction for future investigation. Notably, a recent study reported that NELF depletion is accompanied by transcriptional features associated with aging and cellular senescence [57]. Given the established role of NELF in promoter-proximal pausing [21, 22, 58], this suggests a possible context in which large-scale changes in pausing distribution may be relevant to broader cellular regulatory changes.

Evolutionary variation in sequence composition in the pausing region may also be associated with divergence in promoter-proximal pausing dynamics. Across species comparisons, non-conserved TSS genes showing significantly lower pause-escape rates in human and rhesus macaque compared with baboon also tended to exhibit higher GC content within pausing regions than their baboon orthologs. These observations are consistent with previous models proposing that GC-rich promoter environments may slow early transcription elongation through increased local DNA duplex stability [25, 59]. Pausing dispersion also showed a distinct relationship with GC skew. Broad-to-sharp changes tended to be associated with increased GC skew, whereas sharp-to-broad changes were more frequently associated with reduced GC skew, particularly at non-conserved TSS genes. Previous studies have linked promoter GC skew to R-loop formation [6,43,60], raising the possibility that GC-skew-dependent R-loop formation may influence how paused polymerase accumulates or redistributes near promoters, potentially contributing to variation in pause-site distribution.

Taken together, our results provide a quantitative framework for describing promoter-proximal pausing in terms of kinetic and distributional aspects with distinct patterns of variation and association. This framework makes it possible to compare pausing behavior across biological contexts and to relate these measurements to promoter chromatin and sequence features. Additional work will be required to determine which molecular processes contribute to changes in pause-site organization and pause-escape kinetics, and under what conditions these two aspects of pausing become more tightly coupled.

## Methods

### Likelihood ratio test

We adopt the probabilistic framework previously developed [24, 26] to model promoter-proximal transcriptional pausing under steady-state conditions. In this framework, the read count at each nucleotide position is modeled as a Poisson-distributed variable with mean proportional to local RNA polymerase occupancy. To account for heterogeneity in pause-site location across cells, we introduce a latent variable formulation. For each genomic position *k* ∈ [*k*_min_*, k*_max_], we assume that the fraction of cells with pause sites at position *k*, denoted by *f_k_*, follows a Gaussian distribution parameterized by mean *µ* and dispersion *σ*. Let *Y_k_* denote the (unobserved) number of reads originating from this subpopulation, and *X_k_* the observed read count. The total number of pause-associated reads is *t* = Σ*_k_ Y_k_*. Because *Y*_k_ is not directly observed, we estimate its posterior expectation ⟨*Y_k_*⟩ using an expectation–maximization (EM) algorithm, with ⟨*t*⟩ = Σ*_k_* ⟨*Y_k_*⟩. The Gaussian parameters defining *f_k_* are updated during the M-step. Under this model, the complete-data log-likelihood for a single transcription unit is:

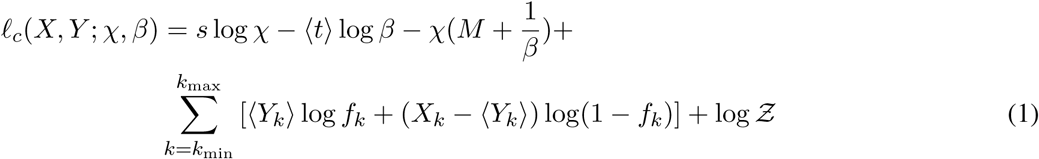

where *s* = Σ*_i_ X_i_* is the total read count, *χ* is estimated as the average read count per base in a downstream region of length *M*, and Z is a normalization constant that does not depend on the free parameters. Terms that are constant across hypotheses (including *s* log *χ*, −*χM*, and log *Z*) cancel in likelihood ratio tests. After cancellation of terms that are constant across hypotheses, the likelihood ratio test statistic depends only on the following component:

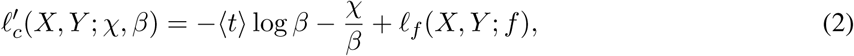

with

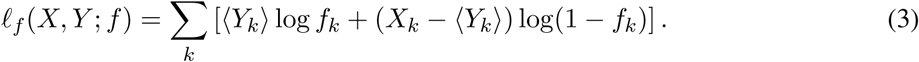

Parameters are estimated using the EM algorithm. In the E-step, ⟨*Y_k_*⟩ and ⟨*t*⟩ are computed. In the M-step, *β* is updated in closed form, while *f_k_* is modeled as a truncated Gaussian over [*k*_min_*, k*_max_] with parameters (*µ, σ*^2^). To compare transcription dynamics between two conditions, we perform likelihood ratio tests (LRTs) for both the pause-escape rate *β* and the pause-site distribution *f_k_*. Under *H*_1_, both *β* and *f_k_* are allowed to differ between conditions. Parameters are estimated independently for each condition, yielding (*β*^^(1)^*, f* ^(1)^) and (*β*^^(2)^*, f* ^(2)^). The joint log-likelihood component is:

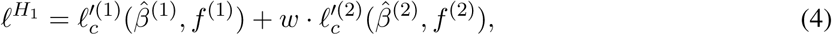

where *w* accounts for differences in sequencing depth. To test for differences in pause-escape kinetics, we constrain *β*^(1)^ = *β*^(2)^ = *β*, while allowing *f_k_*to vary independently. The shared estimate is:

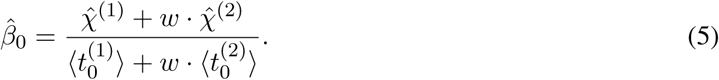

The corresponding LRT-relevant component of the log-likelihood is:

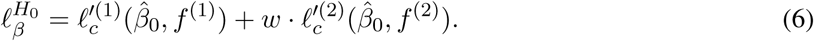

To test for differences in pause-site distribution, we constrain *f* ^(1)^ = *f* ^(2)^ = *f*, while allowing *β* to vary independently. Counts from both conditions are aggregated:

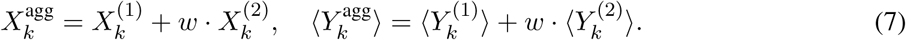

The shared distribution *f_k_* is then estimated from aggregated data, yielding:

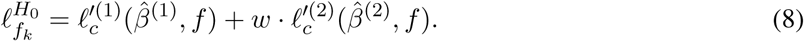

The likelihood ratio statistics are defined as:

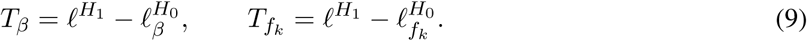

For each gene, 2*T_β_* and 2*T_f__k_* are approximately *χ*^2^-distributed with one and two degrees of freedom, respectively. *p*-values are obtained from these distributions and adjusted using the Benjamini–Hochberg procedure. Conventional *χ*^2^ tests applied to promoter-proximal versus gene-body read counts effectively evaluate the inverse of the pause-escape rate (i.e., the pausing index). The LRT therefore provides a direct statistical test of changes in *β*, while also enabling explicit testing of pause-site distributions via *f_k_*.

Statistical analyses, including parameter estimation and likelihood-ratio tests, were performed using a combination of custom scripts and the STADyUM R package, which provides a user-friendly implementation of the probabilistic framework described here.

### Synthetic NRS data for testing model performance

Synthetic nascent RNA sequencing (NRS) data were generated using the SimPol simulator [24]. Briefly, transcriptional initiation (*αζ*) and pause-escape (*βζ*) rates were varied from 0.1 to 10 events per minute per cell. Elongation rates at each nucleotide position were sampled from a truncated normal distribution (mean = 2,000 nt/min, standard deviation = 1,000 nt/min, truncated to [1,500, 2,500] nt/min). Pause-site distributions were systematically varied by changing the mean (*µ*) and standard deviation (*σ*) of the truncated normal distribution, and bounded [17, 200] nt. Specifically, *µ* was set to 30, 40, 45, 50, 55, 60, 65, 70, 75, 80, and 90 nt, and *σ* was set to 10, 20, 25, 30, 35, 40, 45, 50, 55, 60, and 70 nt, resulting in a grid of parameter combinations.

A fixed center-to-center RNA polymerase spacing of *s_p_* = 50 nt was assumed. For each parameter combination, 20,000 cells were simulated over 40 minutes (400,000 time steps). From each simulation, 5,000 cells were randomly sampled and this sampling was repeated 50 times to generate independent replicates. For each replicate, read counts at position *i* were drawn from a Poisson distribution with mean:

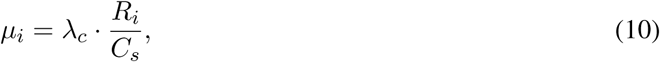

where *R_i_*denotes the number of RNA polymerases at position *i*, *C_s_* = 5, 000 is the number of sampled cells, and *λ_c_*is a scaling factor controlling sequencing depth. The parameter *λ_c_* was calibrated such that simulations with *αζ* = *βζ* = 1 produced gene-body read depths consistent with empirical PRO-seq data [24, 61]. Simulated genes were 2,000 bp in length, with pausing analysis restricted to the first 200 bp. Gene-body read counts were sampled separately using:

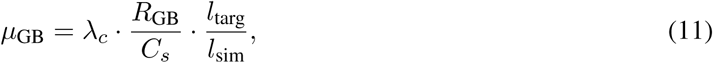

where *l*_targ_ = 19, 800 bp and *l*_sim_ = 1, 800 bp.

### Application of the LRT to synthetic data

For each simulated dataset, likelihood ratio tests were applied to detect differences in pause-escape rates (*β*) and pause-site distributions (*f_k_*) between conditions. Statistical power was evaluated across a range of parameter perturbations, including changes in *β*, *µ*, and *σ*.

### Application of the LRT to experimental datasets

We analyzed PRO-seq datasets from P-TEFb inhibition and NELF depletion experiments reported in ref. [21]. Raw sequencing data were processed using the proseq2.0 pipeline in single-end mode [62]. Reads were mapped to the human genome (GRCh38.p13), and gene annotations were obtained from Ensembl release 99 [63]. Only protein-coding genes located on autosomes and sex chromosomes were retained, and overlapping genes on the same strand were excluded. To define transcription start sites (TSSs), we used PRO-cap data from the same study. PRO-cap signals were lifted over from GRCh37 to GRCh38 and used to refine annotated TSS positions. For each gene, the dominant TSS was defined as the position with the highest PRO-cap signal within 250 bp of the annotated TSS. Promoters with fewer than 10 PRO-cap reads were excluded. All downstream analyses were performed relative to this refined TSS.

For each retained gene, the pause region used for model fitting was defined as the first 200 bp downstream of the annotated TSS. The gene body counting region excluded the 1,250 bp TSS-proximal region and the 1,250 bp transcript-end-proximal region, and was capped at 90 kb. Genes with gene body regions shorter than 6 kb were removed, and genes with 20 or fewer reads in either the pause region or the gene body were excluded. Read counts were summarized as follows: per-base PRO-seq 3*^′^*-end counts were extracted across the 200 bp pause window, and total read counts were aggregated across the gene body counting region. These quantities were used as inputs to the probabilistic pause-release model to estimate pause-escape rates and downstream likelihood ratio statistics.

### Primate PRO-seq data generation and processing

PRO-seq libraries were generated as described previously with minor modifications [4, 64, 65]. Briefly, nuclei were isolated and subjected to nuclear run-on in the presence of biotin-labeled nucleotides to capture engaged RNA polymerases. The run-on reaction was performed in the presence of Sarkosyl to prevent new transcription initiation while allowing engaged RNA polymerases to resume elongation. Nascent RNA was purified and fragmented by base hydrolysis, followed by 3*^′^* adapter ligation. Biotin-labeled RNA was enriched using streptavidin magnetic beads, and 5*^′^* ends were prepared by enzymatic decapping and phosphorylation prior to 5*^′^* adapter ligation. Reverse transcription was performed on bead-bound RNA, and library amplification cycles were determined by qPCR. Libraries were sequenced on Illumina platforms using adapter configurations designed to recover the 3*^′^* ends of nascent RNA, from which RNA polymerase active sites were inferred.

All PRO-seq datasets were processed using proseq2.0 [62] in paired-end mode. Reads were aligned to the reference genomes (human: GRCh38.p13; rhesus: Mmul 10; baboon: Panu 3.0), and the 3*^′^* ends of aligned reads were used to represent RNA polymerase active sites. Only uniquely mapped reads were retained, and strand-specific bigWig files were generated for downstream analyses. Quality control steps included removal of low-quality reads and filtering of genes with insufficient coverage in either promoter-proximal or gene-body regions, as described above. For cross-condition comparisons, analyses were restricted to genes with matched TSS definitions across conditions.

### Definition of pausing dispersion groups

Given that pausing dispersion (*σ*) exhibited a bimodal distribution within cell types (**Fig. 3D** and **Supplementary Fig. S5A**), genes were classified into *Sharp* and *Broad* pausing groups using a data-driven threshold defined by the local minimum between the two modes. Specifically, a histogram of *σ* values was constructed, and the histogram counts were smoothed using a centered moving average with a window size of five bins. Local minima within the bimodal interval (20 *< σ <* 50) were identified using the `findpeaks` function from the `pracma` package applied to the negative smoothed histogram counts, and the minimum with the lowest smoothed count was selected as the threshold. Genes with *σ* values below this threshold were classified as *Sharp* pausing, and genes with *σ* values above this threshold were classified as *Broad* pausing.

### Micro-C data generation and 1D signal processing

Micro-C experiments were performed as described previously [16]. Briefly, cells were subjected to dual crosslinking using formaldehyde followed by disuccinimidyl glutarate (DSG) to stabilize chromatin interactions. Chromatin was digested with micrococcal nuclease (MNase) to achieve predominantly mononucleosome fragments, followed by end repair, biotin fill-in, and in situ proximity ligation. After ligation, biotin-labeled DNA fragments were enriched using streptavidin beads and processed into sequencing libraries using standard protocols. Libraries were sequenced in paired-end mode on Illumina platforms.

Sequencing reads were mapped to the reference genome and filtered for high-quality intrachromosomal contacts as previously described [16]. Contact maps were generated and normalized for quality control. Reads were further filtered to retain pairs with mapping quality (MAPQ) ≥ 30 on both ends. For downstream analyses, Micro-C signals were converted into one-dimensional genomic profiles by projecting read positions onto genomic coordinates and shifting each read end by 75 bp according to strand orientation to approximate nucleosome centers [16]. For each gene, these signals were extracted within ±1 kb windows centered on refined TSSs using `PyRanges`, and aggregated to generate per-base coverage profiles. The summed Micro-C signal within the 0–200 bp region downstream of the TSS was used as a measure of promoter-proximal nucleosome occupancy. These one-dimensional signals were further used to quantify promoter-proximal positioning as described below.

### Histone modification analysis

Histone modification bigWig files for human CD4^+^ T cells and CD14^+^ monocytes were obtained from ENCODE [37]. For each gene, signal intensity was quantified as the mean bigWig signal within ±500 bp windows centered on refined TSSs. To ensure comparability across cell types, analyses were restricted to genes with matched TSS definitions. Differences in histone modification signals between cell types were quantified using DiffBind [66] applied to these shared promoter regions.

### Promoter sequence analysis

Core promoter annotations were obtained from the Eukaryotic Promoter Database (EPD) [67]. For each refined TSS, the nearest annotated EPD promoter within 200 bp on the same strand and gene was assigned. Promoters were classified into motif categories based on the presence of core elements, including TATA-box, Inr, GC-box, and CCAAT-box. Categories included TATA-box, GC–CCAAT, CCAAT-only, GC-only, GC–Inr, Inr-only, and Other.

For each gene expressed in human (hg38), rhesus macaque (rheMac10), and olive baboon (papAnu4) CD4^+^ T cells and CD14^+^ monocytes, TSS-centered FASTA sequences were extracted from the corresponding reference genomes using `bedtools getfasta`. GC content and GC skew were calculated using a 100 bp sliding window across the extracted regions, and average profiles were visualized as metaplots relative to the TSS. Gene-level GC content and GC skew were additionally summarized using the 200 bp region immediately downstream of the TSS. Delta GC content was defined as the log_2_ fold change between orthologous gene pairs, whereas delta GC skew was defined as the arithmetic difference.

### Gene ontology analysis

Genes were grouped according to pause-escape rate categories, and Gene Ontology (GO) biological process enrichment analysis was performed using `clusterProfiler` [68]. Enrichment was computed using `enrichGO` with all genes exhibiting detectable transcription signals as background. Redundant GO terms were reduced using the `simplify` function with a semantic similarity cutoff of 0.9.

### Quantification of nucleosome positioning

Nucleosome positioning was quantified using Micro-C signal within the 200 bp region downstream of the TSS. For each gene, the signal was normalized to a probability distribution across positions, and Shannon entropy was computed as:

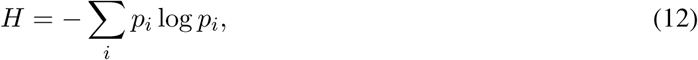

where *p_i_* is the normalized signal at position *i*. Entropy was normalized by its theoretical maximum (log *L*, with *L* = 200) to obtain values between 0 and 1. Lower entropy indicates more localized nucleosome positioning, whereas higher entropy indicates more dispersed signal. To facilitate interpretation, nucleosome positioning strength was defined as (1 − *H*). Differences between conditions were quantified as:

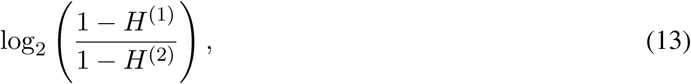

where *H*^(1)^ and *H*^(2)^ denote normalized entropy values in the two conditions.

### Identification of conserved and non-conserved TSSs

One-to-one orthologous genes between species were identified using OrthoFinder [69]. Promoter-proximal regions were mapped to the human reference genome using UCSC liftOver [70]. For each ortholog pair, overlap between promoter-proximal pausing regions was computed. Genes with overlap greater than 150 bp were classified as having conserved TSSs, whereas those with overlap less than 150 bp were classified as non-conserved TSSs.

## Data availability

Published data were retrieved from GEO and ENCODE (see **Supplementary Table S1–2** for details), including perturbation PRO-seq data from Gene Expression Omnibus (GEO) accession GSE144786. Newly generated PRO-seq and Micro-C data have been deposited in dbGaP under accession phs002146.v2 and in the GEO under accession GSE342012 and GSE342013 (see **Supplementary Tables S3–S4**). Processed data files used in this study are available from Zenodo (doi.org/10.5281/zenodo.20598895).

## Code availability

The STADyUM package is available through Bioconductor (https://bioconductor.org/packages/STADyUM). All code required to reproduce the results presented in this study is available from the GitHub repository https://github.com/EasyPiPi/unimod%5Fpausing.

## Author contributions

Xin Zeng: Formal analysis (lead); Software (supporting); Visualization (lead); Writing – original draft (lead); Writing – review and editing (supporting). Gilad Barshad: Investigation (lead); Resources (supporting). Rebecca Hassett: Software (lead). Edward J. Rice: Investigation (lead); Resources (supporting). Charles G. Danko: Investigation (supporting); Resources (lead); Funding acquisition (lead); Writing – review and editing (supporting). Adam Siepel: Conceptualization (lead); Methodology (lead); Resources (supporting); Funding acquisition (lead); Supervision (lead); Writing – review and editing (lead). Yixin Zhao: Conceptualization (lead); Methodology (lead); Software (lead); Formal analysis (supporting); Supervision (supporting); Writing – original draft (lead); Writing – review and editing (supporting).

## Acknowledgements

We thank members of the Siepel and Danko laboratories for helpful discussions. This study is supported by National Institute of General Medical Sciences (R35-GM127070 to A.S.) and National Human Genome Research Institute (R01-HG012944 to A.S. and R01-HG010346 to C.G.D.). The content is solely the responsibility of the authors and does not necessarily represent the official views of the US National Institutes of Health.

