## supplementary figures and tables for "A comparative analysis of promoter-proximal pausing reveals kinetic and distributional dimensions of variation"

### Supplementary Information for:
### A comparative analysis of promoter-proximal pausing reveals kinetic and distributional dimensions of variation

Xin Zeng[1], Gilad Barshad[2], Rebecca Hassett[1], Edward J Rice[2],
Charles G Danko[2], Adam Siepel[1,*], and Yixin Zhao[1,3,*]

[1]Simons Center for Quantitative Biology, Cold Spring Harbor Laboratory, Cold Spring Harbor, NY 11724, USA
[2]Baker Institute for Animal Health, College of Veterinary Medicine, Cornell University, Ithaca, NY 14853, USA
[3]Yazhouwan National Laboratory, Sanya, Hainan 572025, China

### Supplementary Figures

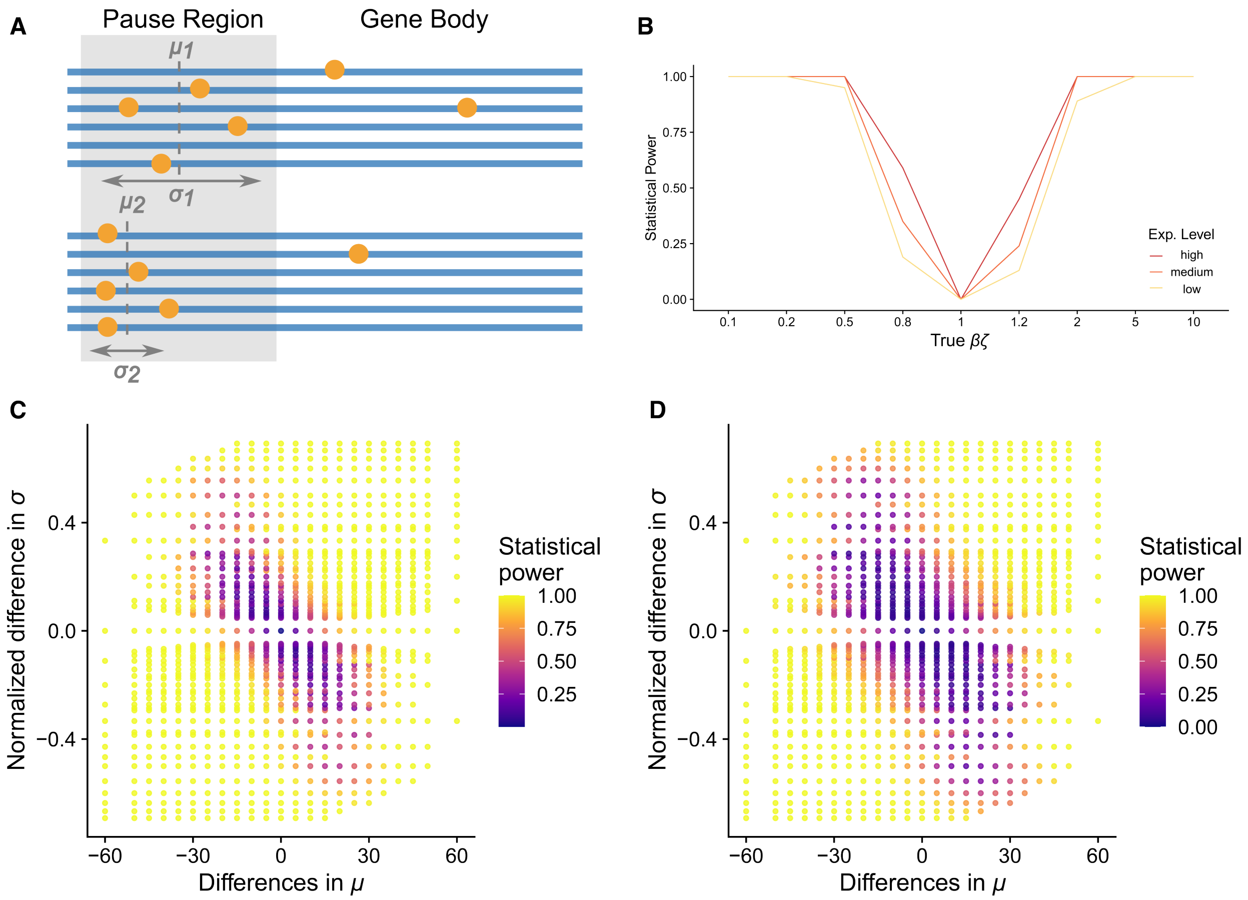


Supplementary Figure S1: Likelihood-ratio testing framework evaluated using simulated nascent RNA profiles. **A.** Simulation scheme for generating nascent RNA profiles. RNA polymerase (RNAP) initiation and pause-escape are modeled using specified rate parameters, and pause positions are sampled from Gaussian distributions parameterized by the mean ($\mu$) and standard deviation ($\sigma$). Reads are drawn from the resulting RNAP density across promoter-proximal and gene-body regions. **B.** Statistical power of the LRT for detecting differences in pause-escape rates across high-, medium-, and low-expression genes. **C–D.** Two-dimensional power surfaces of the distributional LRT for detecting coordinated shifts in pause-site mean ($\mu$) and dispersion ($\sigma$), shown for **(C)** medium- and **(D)** low-expression genes.

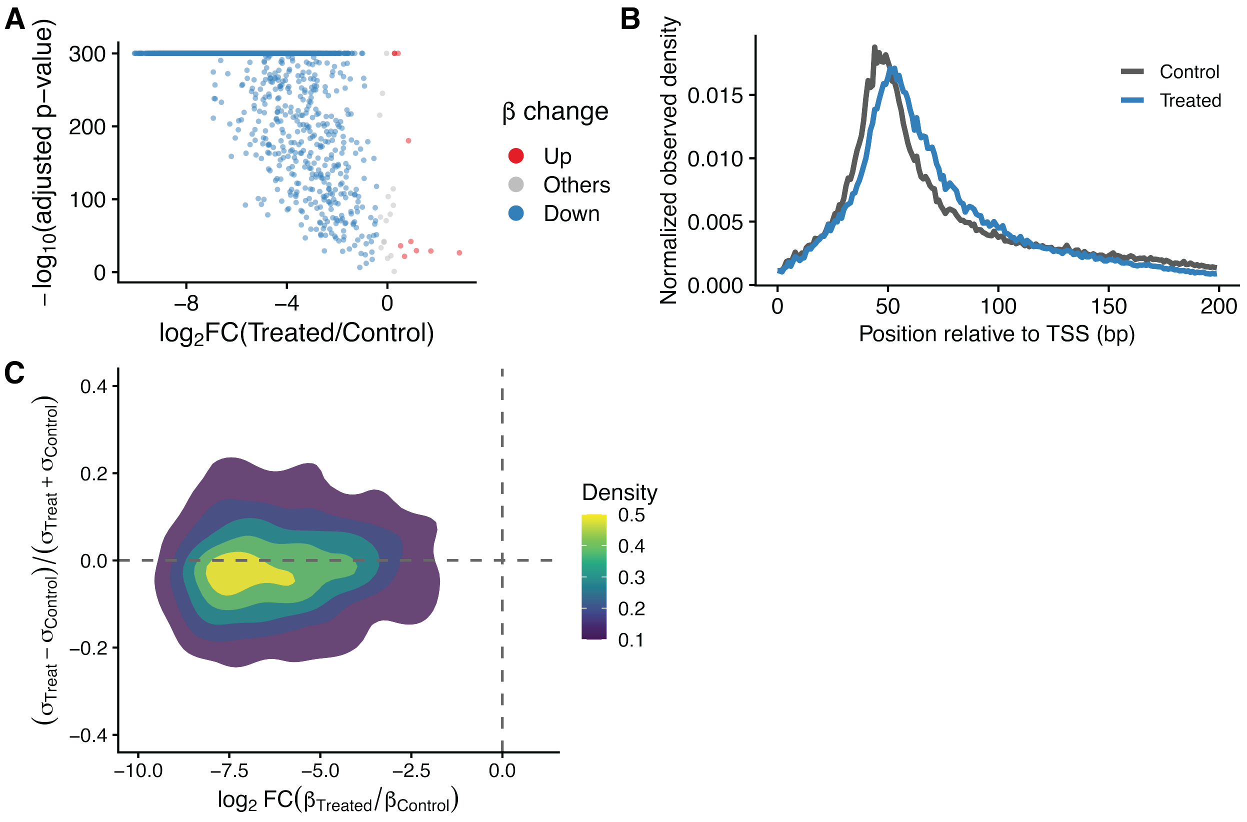


Supplementary Figure S2: Perturbation analyses following flavopiridol treatment. **A.** Genome-wide LRT results for pause-escape rates ($\beta$) following flavopiridol treatment. **B.** Metaplots of normalized PRO-seq density downstream of TSS in control and flavopiridol-treated cells. **C.** Global distributions of $\log_2$ fold changes in $\beta$ and pausing dispersion ($\sigma$) following flavopiridol treatment.

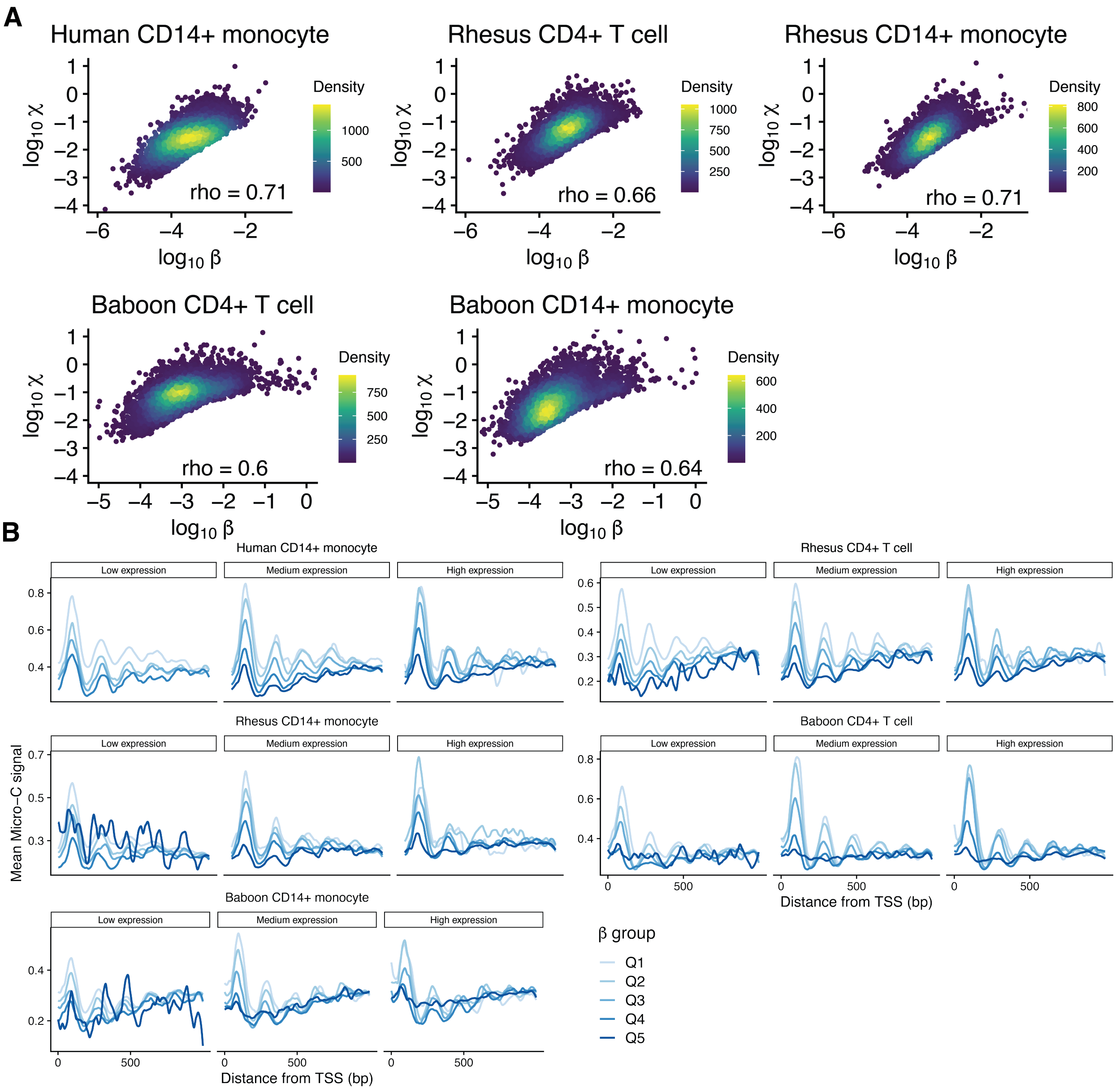


Supplementary Figure S3: Relationships among pause-escape kinetics, transcriptional activity, and promoter chromatin structure within cell types. **A.** Relationship between $\beta$ and $\chi$ across species and cell types. Scatter plots show $\log_{10}$-transformed $\beta$ and $\chi$ values for individual genes. Point color indicates local density, and Spearman correlation coefficients are shown. **B.** Average Micro-C signal around TSSs is shown for genes stratified by $\beta$ quintiles (Q1–Q5) and further grouped by $\chi$ into low, medium, and high categories.

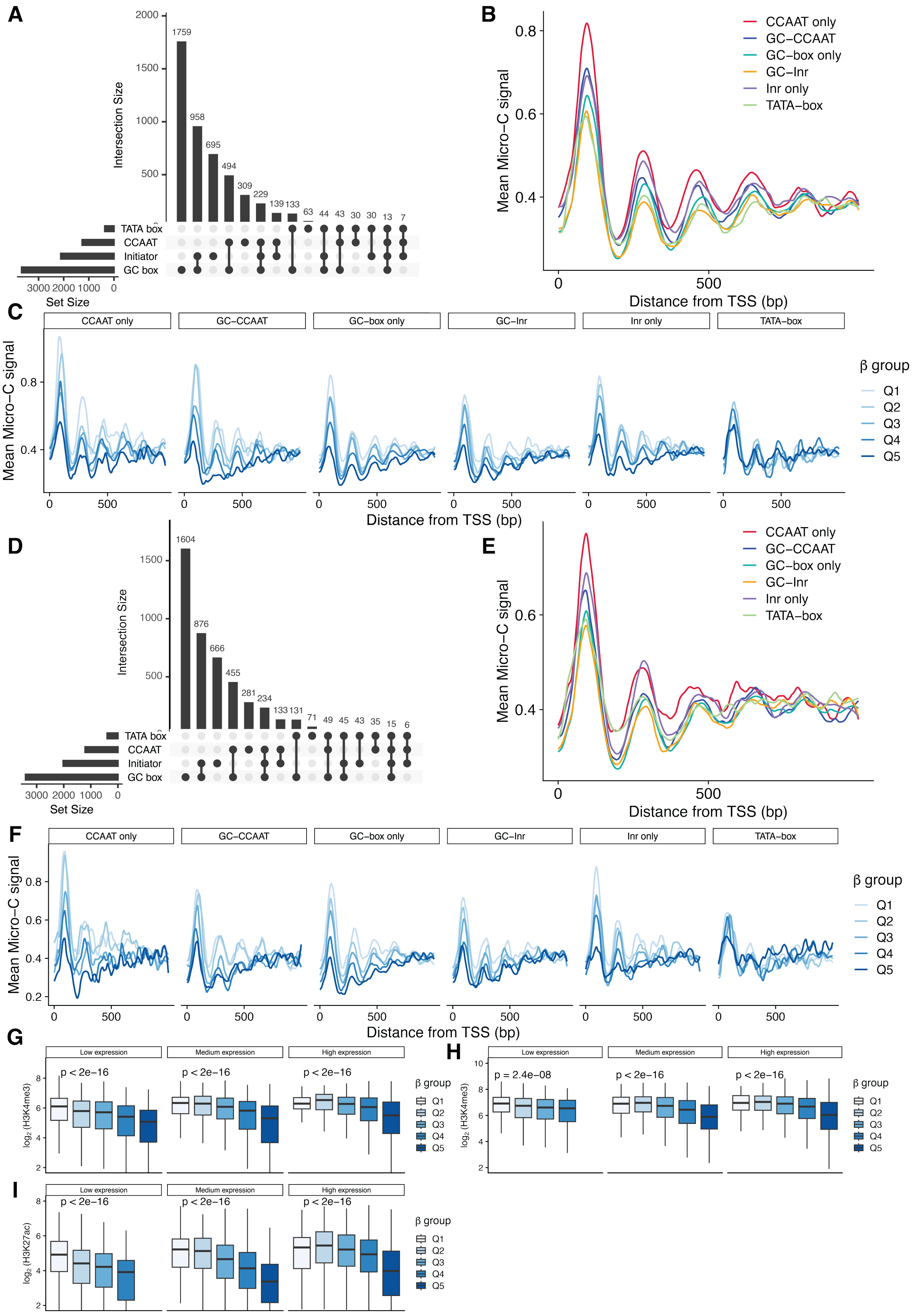


Supplementary Figure S4

**Supplementary Figure S4 continued.**

Promoter architecture, nucleosome organization, and chromatin features across human $CD4^+$ T cells and $CD14^+$ monocytes. **A–D.** UpSet plots of promoter motif combinations among expressed genes in human $CD4^+$ T cells (**A**) and $CD14^+$ monocytes (**D**). **B–E.** Average Micro-C signal around TSSs for genes grouped by promoter motif class in human $CD4^+$ T cells (**B**) and $CD14^+$ monocytes (**E**). **C–F.** Average Micro-C signal around TSSs for genes in different promoter motif classes, stratified by $\beta$ quintiles (Q1–Q5), in human $CD4^+$ T cells (**C**) and $CD14^+$ monocytes (**F**). **G–H.** Boxplots showing H3K4me3 levels for genes stratified by $\beta$ quintiles within low, medium, and high expression groups in human $CD4^+$ T cells (**G**) and $CD14^+$ monocytes (**H**). **I.** Boxplots showing H3K27ac levels for genes stratified by $\beta$ quintiles within low, medium, and high expression groups in human $CD14^+$ monocytes. For panels G–I, P values were calculated using Kruskal–Wallis tests across the five $\beta$ groups.

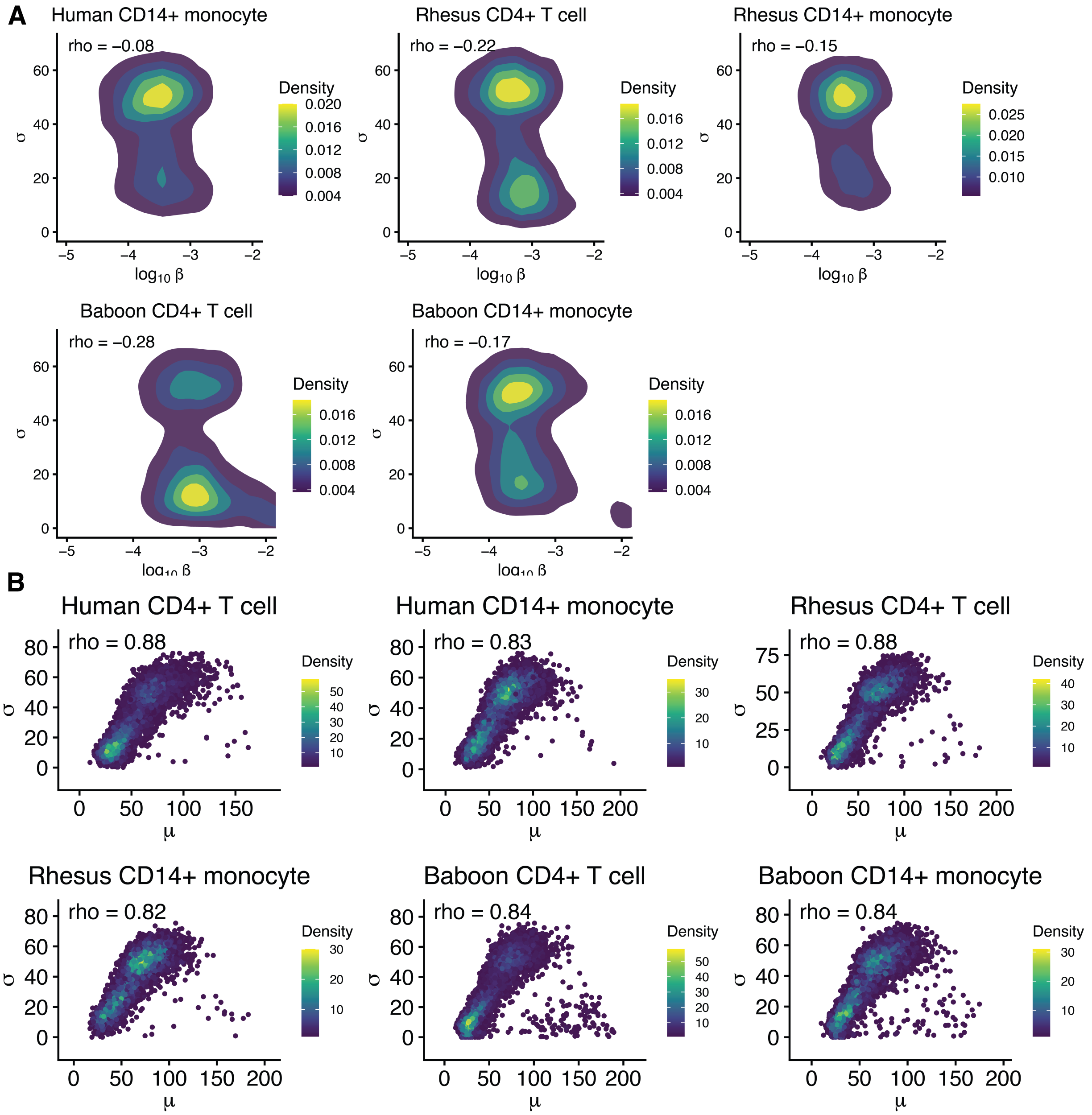


Supplementary Figure S5: Relationships among promoter-proximal pausing parameters within cell types. **A.** Relationship between $\beta$ and pausing dispersion ($\sigma$). Contour plots show the density distribution of genes by $\beta$ and $\sigma$. Spearman correlation coefficients are shown. **B.** Relationship between mean pause position ($\mu$) and pausing dispersion ($\sigma$). Scatter plots show individual genes colored by local density. Spearman correlation coefficients are shown.

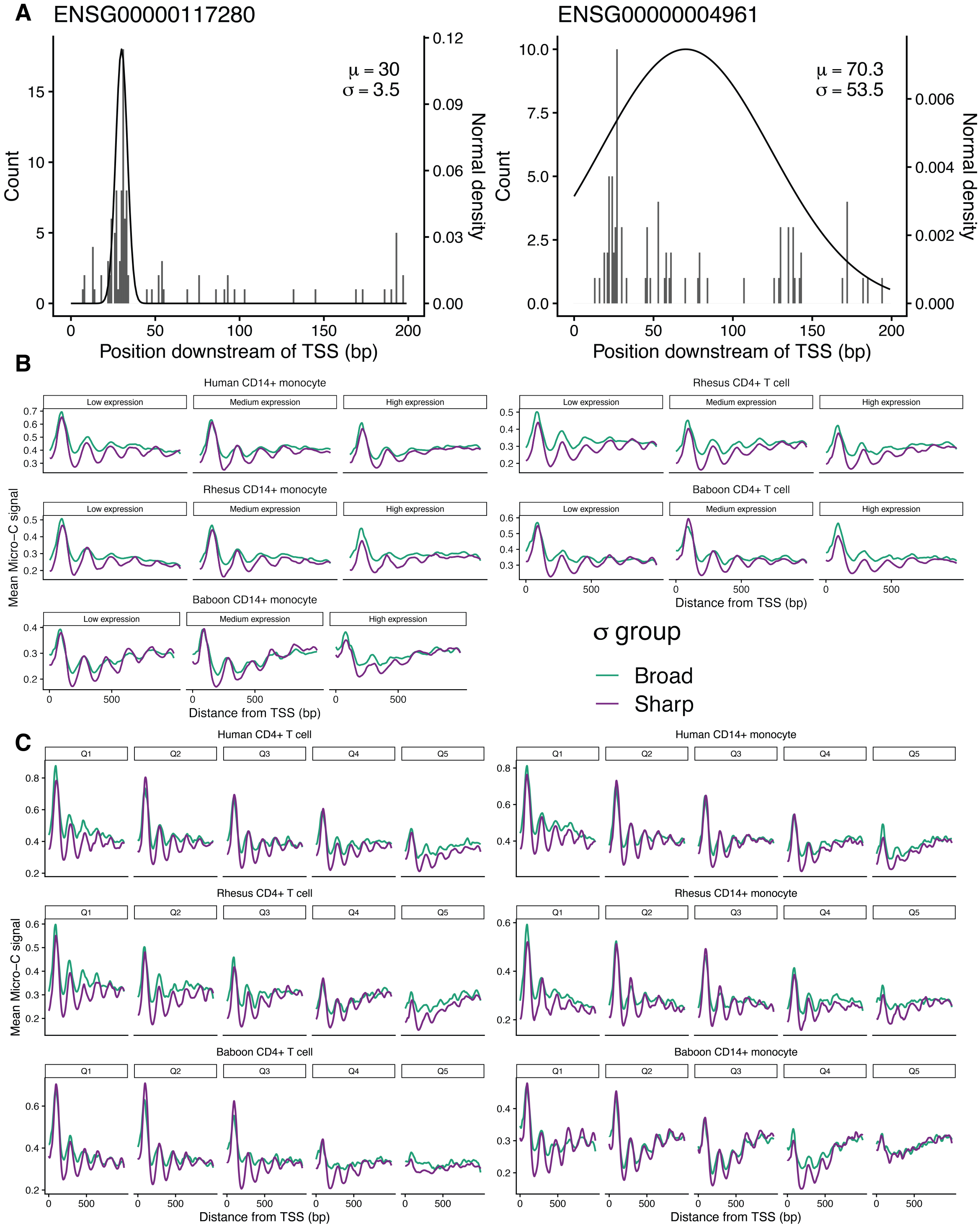


Supplementary Figure S6: Pausing dispersion is associated with distinct promoter chromatin organization. **A.** Representative genes illustrating sharp and broad pausing patterns are shown. For each gene, PRO-seq signal downstream of the TSS is displayed together with a fitted distribution (black curve), with corresponding mean and dispersion ($\sigma$). **B.** Average Micro-C signal around TSSs is shown for genes stratified by pausing dispersion ($\sigma$) groups, with genes classified as broad or sharp pausing, and further grouped by $\chi$ into low, medium, and high categories. **C.** Average Micro-C signal around TSSs is shown for genes stratified by $\beta$ quintiles (Q1–Q5), with genes further grouped by pausing dispersion ($\sigma$) as broad or sharp pausing.

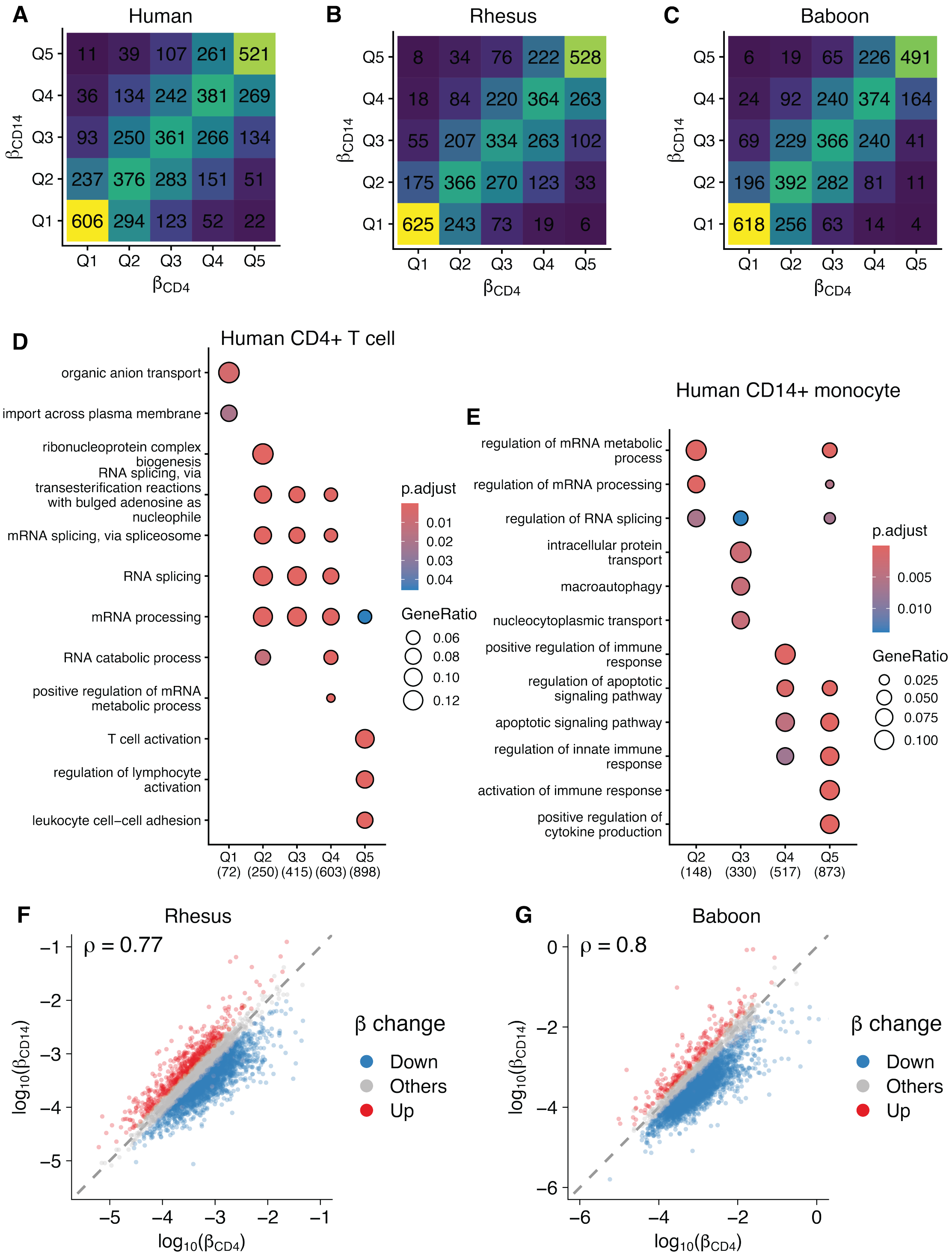


Supplementary Figure S7: Comparative analysis of pause-escape kinetics across cell types within species. **A–C.** Global ordering of pause-escape kinetics is preserved across cell types in human (**A**), rhesus macaque (**B**), and olive baboon (**C**). Genes are stratified into $\beta$ quintiles (Q1–Q5) in each cell type, and heatmaps show the number of genes in each pairwise quintile bin. **D.** Gene Ontology (GO) enrichment analysis for genes in different $\beta$ groups within the high $\chi$ group in human CD4$^+$ T cells. **E.** Same analysis within the high $\chi$ group for human CD14$^+$ monocytes. Dot size indicates GeneRatio, and color indicates adjusted $p$ value (see **Methods**). **F–G.** Comparison of pause-escape rates ($\beta$) between CD4$^+$ T cells and CD14$^+$ monocytes in rhesus macaque (**F**) and olive baboon (**G**). Genes are colored according to LRT-inferred significant changes in $\beta$.

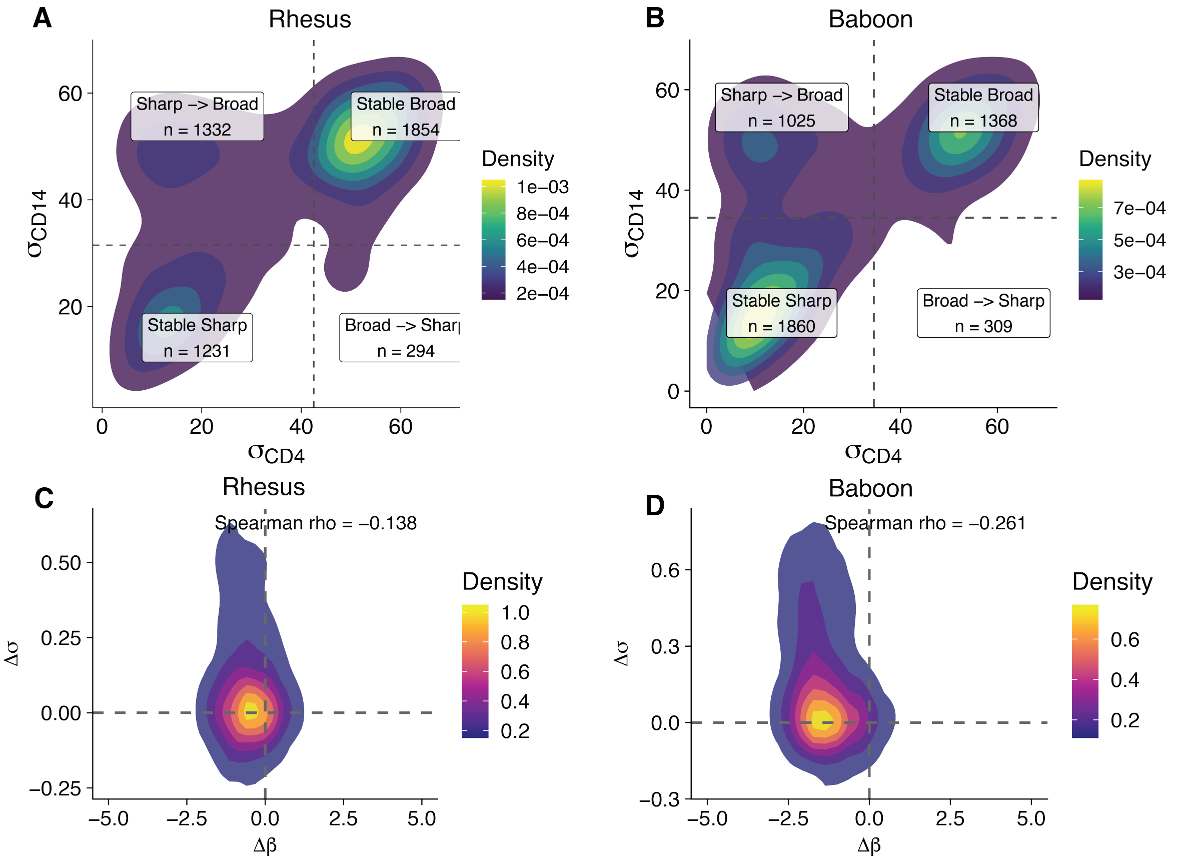


Supplementary Figure S8: Comparative analysis of pausing dispersion and pause-escape kinetics across cell types within species. **A–B.** Comparison of pausing dispersion ($\sigma$) between CD4$^+$ T cells and CD14$^+$ monocytes within rhesus macaque (**A**) and olive baboon (**B**). Genes are classified according to broad and sharp pausing dispersion groups in each cell type. **C–D.** Joint distribution of changes in pause-escape rate ($\Delta\beta$) and pausing dispersion ($\Delta\sigma$) across cell types within rhesus macaque (**C**) and olive baboon (**D**). Spearman correlation coefficients are shown.

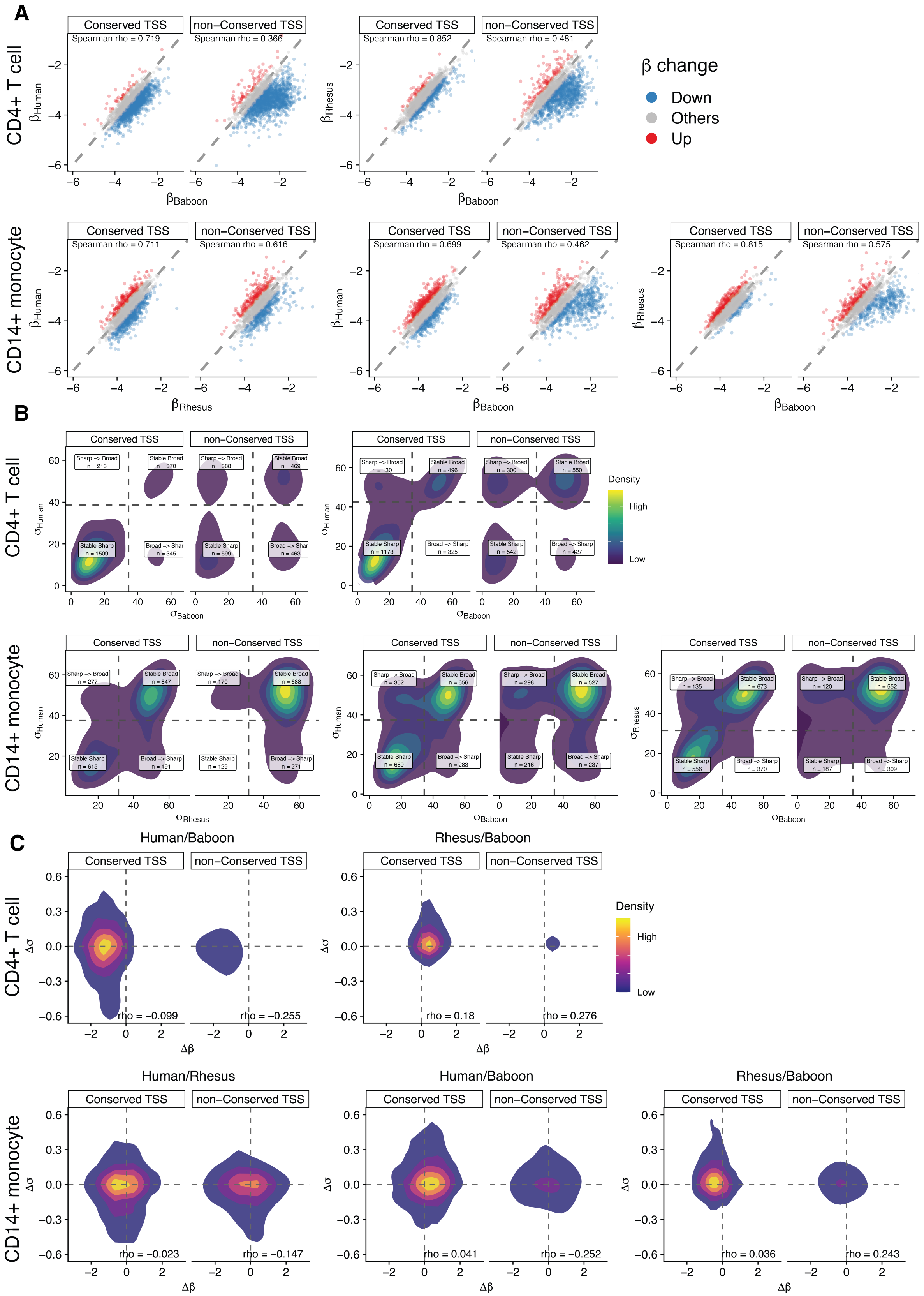


Supplementary Figure S9

**Supplementary Figure S9 continued.**

Cross-species comparison of pause-escape rate and pausing dispersion. **A.** Comparison of pause-escape rates ($\beta$) between orthologous genes across species. Conserved and non-conserved TSS genes are shown separately. Genes are colored according to LRT-inferred significant changes in $\beta$. Spearman correlation coefficients are shown. **B.** Comparison of pausing dispersion groups ($\sigma$) between orthologous genes across species. Orthologous genes are classified into stable sharp, stable broad, sharp-to-broad, and broad-to-sharp categories based on pausing dispersion groups in each species. Conserved and non-conserved TSS genes are shown separately. Density contours represent the distribution of genes by pausing dispersion. **C.** Joint distribution of changes in pause-escape rate ($\Delta\beta$) and pausing dispersion ($\Delta\sigma$) across species. Density contours represent the distribution of orthologous genes by $\Delta\beta$ and $\Delta\sigma$. Conserved and non-conserved TSS genes are shown separately. Spearman correlation coefficients are shown.

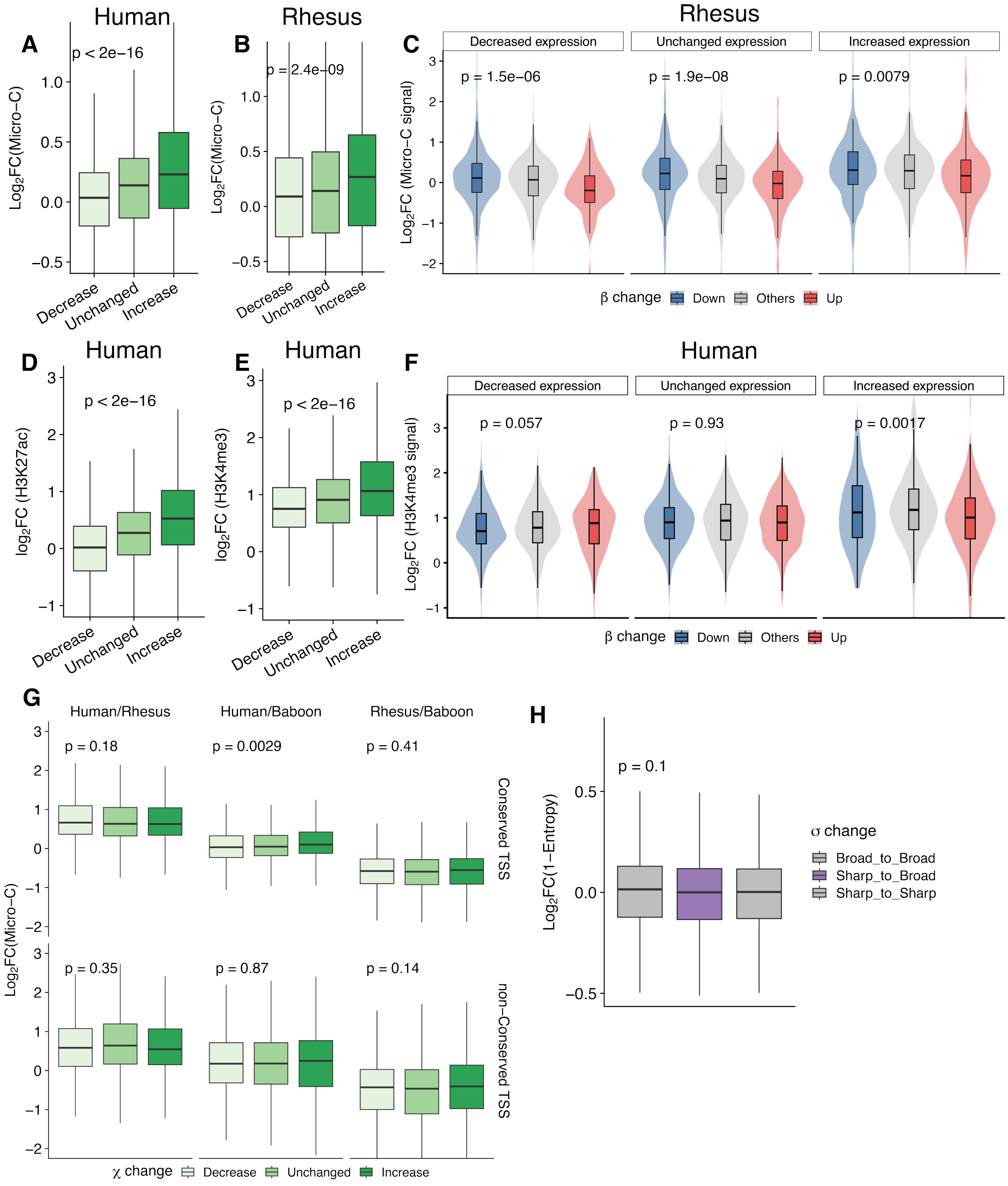


Supplementary Figure S10

**Supplementary Figure S10 continued.**

Chromatin features associated with transcriptional activity ($\chi$), pause-escape rate ($\beta$), and pausing dispersion ($\sigma$) across cell types and species. **A–B.** +1 nucleosome occupancy changes across genes with decreased, unchanged, or increased transcriptional activity ($\chi$) in human (**A**) and rhesus macaque (**B**). **C.** +1 nucleosome occupancy changes in rhesus macaque, stratified by $\chi$ change and direction of pause-escape rate ($\beta$) change. **D–E.** Changes in promoter-associated histone modifications H3K27ac (**D**) and H3K4me3 (**E**) across genes grouped by $\chi$ change in human. **F.** H3K4me3 changes in human, stratified by $\chi$ change and direction of $\beta$ change. **G.** Cross-species changes in +1 nucleosome occupancy across pairwise comparisons in CD4$^{+}$ T cells, grouped by $\chi$ change and TSS conservation status. **H.** Changes in +1 nucleosome positioning, quantified as $1 -$ entropy, across genes grouped by pausing dispersion ($\sigma$) change between cell types. No significant differences are observed across groups. P values were calculated using Kruskal–Wallis tests across the indicated groups.

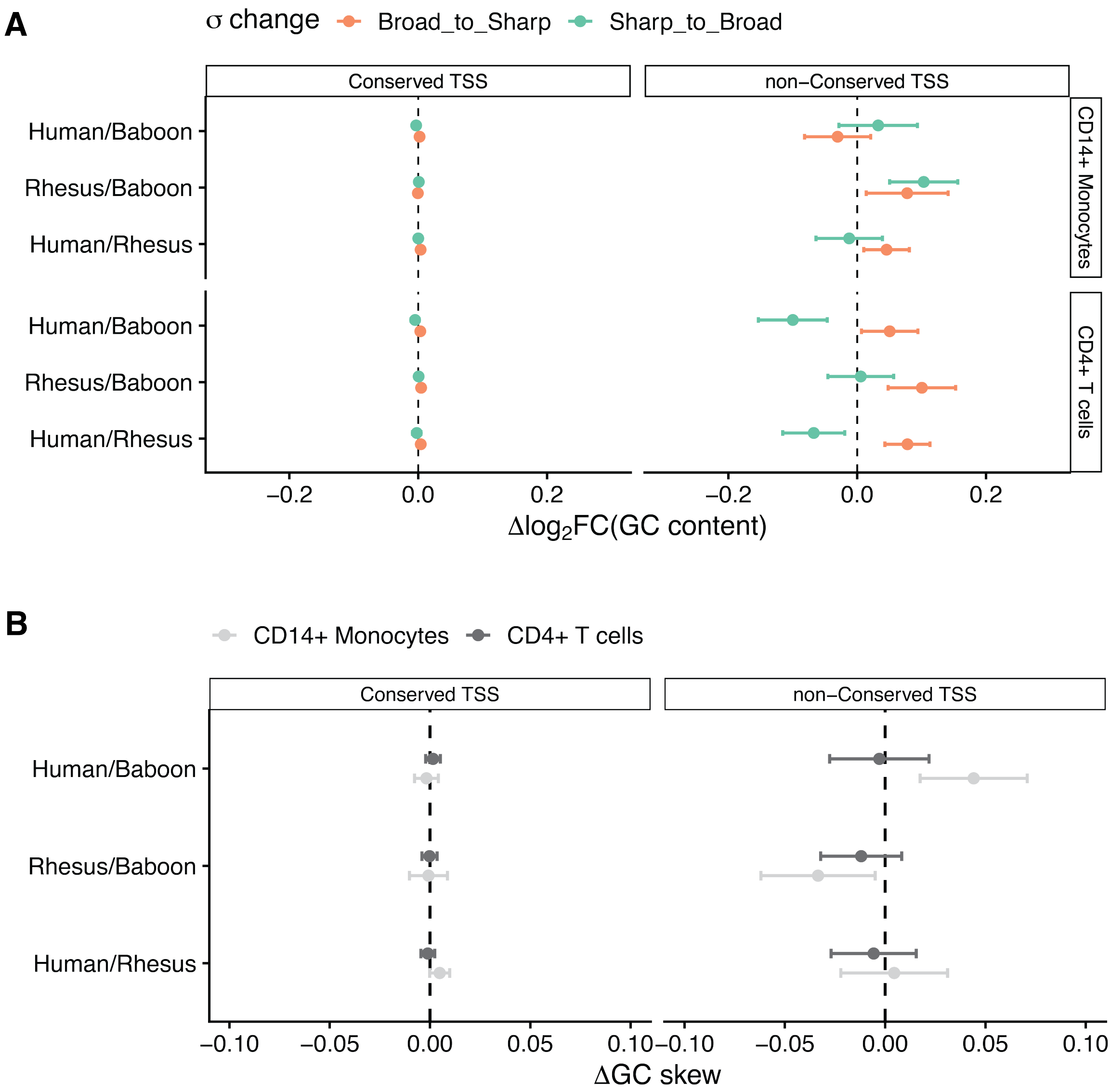


Supplementary Figure S11: Sequence features associated with cross-species variation in pause-escape kinetics and pausing distribution. **A.** Association between cross-species changes in pausing dispersion ($\sigma$) and promoter GC content. Effect sizes are defined as the difference in mean $\log_2$ fold change in GC content between genes with stable and changing pausing distributions. **B.** Association between cross-species differences in pause-escape rate ($\beta$) and GC skew in promoter-proximal regions. Effect sizes are defined as the difference in mean GC skew change between $\beta_{\text{Down}}$ genes and genes with stable $\beta$ ($\beta_{\text{Others}}$) across species and TSS conservation groups.

#### Supplementary Tables

##### Supplementary Table S1. ENCODE ChIP-seq bigwig datasets used for within-cell-type analyses

| accession | donor | histone mark | cell type | experiment accession |
|---|---|---|---|---|
| ENCFF917XJH | ENCDO922TOC | H3K27ac | CD14 monocyte | ENCSR685BBN |
| ENCFF090HMA | ENCDO922TOC | H3K4me3 | CD14 monocyte | ENCSR665QZU |
| ENCFF329UOR | ENCDO922TOC | H3K27ac | CD4 regulatory | ENCSR360IRQ |
| ENCFF811QOJ | ENCDO922TOC | H3K4me3 | CD4 regulatory | ENCSR299XIC |

##### Supplementary Table S2. ENCODE ChIP-seq bam datasets used for between-cell-type analyses

| accession | donor | histone mark | cell type | experiment accession | role |
|---|---|---|---|---|---|
| ENCFF685QTT | ENCDO553CCA | Control | CD14 monocyte | NA | input control |
| ENCFF107RXY | ENCDO836ICQ | Control | CD14 monocyte | NA | input control |
| ENCFF262PYO | ENCDO922TOC | Control | CD14 monocyte | NA | input control |
| ENCFF787FTC | ENCDO553CCA | H3K27ac | CD14 monocyte | ENCSR191YDG | chip |
| ENCFF236PGO | ENCDO836ICQ | H3K27ac | CD14 monocyte | ENCSR319HLH | chip |
| ENCFF516XHC | ENCDO922TOC | H3K27ac | CD14 monocyte | ENCSR685BBN | chip |
| ENCFF018IIB | ENCDO553CCA | H3K4me3 | CD14 monocyte | ENCSR905UNZ | chip |
| ENCFF537WVC | ENCDO836ICQ | H3K4me3 | CD14 monocyte | ENCSR245YME | chip |
| ENCFF228CFW | ENCDO922TOC | H3K4me3 | CD14 monocyte | ENCSR665QZU | chip |
| ENCFF655VRG | ENCDO553CCA | Control | CD4 memory | NA | input control |
| ENCFF512RLD | ENCDO836ICQ | Control | CD4 memory | NA | input control |
| ENCFF991QSO | ENCDO922TOC | Control | CD4 memory | NA | input control |
| ENCFF673UVU | ENCDO553CCA | H3K27ac | CD4 memory | ENCSR832UMM | chip |
| ENCFF538RVA | ENCDO836ICQ | H3K27ac | CD4 memory | ENCSR696DVE | chip |
| ENCFF648XHV | ENCDO922TOC | H3K27ac | CD4 memory | ENCSR175IGC | chip |
| ENCFF071AKA | ENCDO553CCA | H3K4me3 | CD4 memory | ENCSR341QLC | chip |
| ENCFF621MTB | ENCDO836ICQ | H3K4me3 | CD4 memory | ENCSR373MTM | chip |
| ENCFF744HJM | ENCDO922TOC | H3K4me3 | CD4 memory | ENCSR048ARD | chip |

#### Supplementary Table S3. PRO-seq dataset used in this study

| Sample | Species | CellType | Sex | LifeStage | RunType |
|---|---|---|---|---|---|
| HM1_CD4 | HUMAN | CD4 | M | ADULT | PE |
| HM2_CD4 | HUMAN | CD4 | M | ADULT | PE |
| HF1_CD4 | HUMAN | CD4 | F | ADULT | PE |
| HF3_CD4 | HUMAN | CD4 | F | ADULT | PE |
| HM1_CD14 | HUMAN | CD14 | M | ADULT | PE |
| HM2_CD14 | HUMAN | CD14 | M | ADULT | PE |
| HF1_CD14 | HUMAN | CD14 | F | ADULT | PE |
| HF3_CD14 | HUMAN | CD14 | F | ADULT | PE |
| RM1_CD4 | RHESUS | CD4 | M | ADULT | PE |
| RF2_CD4 | RHESUS | CD4 | F | ADULT | PE |
| RM4_CD4 | RHESUS | CD4 | M | ADULT | PE |
| RF4_CD4 | RHESUS | CD4 | F | ADULT | PE |
| RM1_CD14 | RHESUS | CD14 | M | ADULT | PE |
| RF2_CD14 | RHESUS | CD14 | F | ADULT | PE |
| RM4_CD14 | RHESUS | CD14 | M | ADULT | PE |
| RF4_CD14 | RHESUS | CD14 | F | ADULT | PE |
| BM1_CD4 | BABOON | CD4 | M | ADULT | PE |
| BF2_CD4 | BABOON | CD4 | F | ADULT | PE |
| BM3_CD4 | BABOON | CD4 | M | ADULT | PE |
| BM4_CD4 | BABOON | CD4 | M | ADULT | PE |
| BF3_CD4 | BABOON | CD4 | F | ADULT | PE |
| BM5_CD4 | BABOON | CD4 | M | ADULT | PE |
| BF5_CD4 | BABOON | CD4 | F | ADULT | PE |
| BM1_CD14 | BABOON | CD14 | M | ADULT | PE |
| BF2_CD14 | BABOON | CD14 | F | ADULT | PE |
| BM3_CD14 | BABOON | CD14 | M | ADULT | PE |
| BM4_CD14 | BABOON | CD14 | M | ADULT | PE |
| BF3_CD14 | BABOON | CD14 | F | ADULT | PE |
| BM5_CD14 | BABOON | CD14 | M | ADULT | PE |
| BF5_CD14 | BABOON | CD14 | F | ADULT | PE |

**Supplementary Table S4. MicroC dataset used in this study**

| Sample | Species | CellType | Sex | LifeStage |
|---|---|---|---|---|
| HF1_CD4 | HUMAN | CD4 | F | ADULT |
| HM1_CD4 | HUMAN | CD4 | M | ADULT |
| HM2_CD4 | HUMAN | CD4 | M | ADULT |
| HF1_CD14 | HUMAN | CD14 | F | ADULT |
| HM1_CD14 | HUMAN | CD14 | M | ADULT |
| HM2_CD14 | HUMAN | CD14 | M | ADULT |
| RF4_CD4 | RHESUS | CD4 | F | ADULT |
| RF2_CD4 | RHESUS | CD4 | F | ADULT |
| RM4_CD4 | RHESUS | CD4 | M | ADULT |
| RM5_CD14 | RHESUS | CD14 | M | ADULT |
| RF4_CD14 | RHESUS | CD14 | F | ADULT |
| BM5_CD4 | BABOON | CD4 | M | ADULT |
| BF2_CD4 | BABOON | CD4 | F | ADULT |
| BF5_CD4 | BABOON | CD4 | F | ADULT |
| BF2_CD14 | BABOON | CD14 | F | ADULT |
| BM5_CD14 | BABOON | CD14 | M | ADULT |
| BF5_CD14 | BABOON | CD14 | F | ADULT |
| BF3_CD14 | BABOON | CD14 | F | ADULT |
